# Small ARF-like 2 GTPase TITAN 5 is linked with the dynamic regulation of IRON-REGULATED TRANSPORTER 1

**DOI:** 10.1101/2023.04.27.538571

**Authors:** Inga Mohr, Monique Eutebach, Marie C. Knopf, Naima Schommen, Regina Gratz, Kalina Angrand, Lara Genders, Tzvetina Brumbarova, Petra Bauer, Rumen Ivanov

**Affiliations:** Institute of Botany, Heinrich Heine University, Universitätsstr. 1, 40225 Düsseldorf, Germany; former address: Department of Biosciences-Plant Biology, Saarland University, 66123 Saarbrücken, Germany; Cluster of Excellence on Plant Sciences (CEPLAS), Heinrich-Heine University, 40225 Düsseldorf, Germany

**Keywords:** ARF-like, ARLC1, ARL2, EHB1, endomembrane, GTPase, HALLIMASCH, iron, PATL2, plasma membrane, SEC14, SNX1, vesicle

## Abstract

Iron acquisition is crucial for plants. The abundance of IRON-REGULATED TRANSPORTER 1 (IRT1) at the plasma membrane is controlled through endomembrane trafficking. Vesicular trafficking requires small ARF-like GTPases, one of them is TITAN 5 (TTN5). Its physiological functions during the life cycle and cellular targets remain unknown. Little is known how the vesicular trafficking mechanism affects IRT1 localization.

We show that TTN5 interacts with the large cytoplasmic variable region and protein-regulatory platform of IRT1. *ttn5-1^+/-^* plants have a reduced activity of root iron reductase, needed for iron uptake via IRT1. Fluorescent fusion proteins of TTN5 and IRT1 colocalize at the plasma membrane and in endosomes/multivesicular bodies, where IRT1 sorting and cycling between the plasma membrane and the vacuole are coordinated. Colocalization at the plasma membrane depends partly on the interaction ability of TTN5. TTN5 can also interact with peripheral membrane proteins that are components of the IRT1 regulation machinery, like the trafficking factor SNX1, the C2 domain protein EHB1 and the SEC14-GOLD protein PATL2. Hence, this work links iron acquisition and vesicular trafficking involving a small GTPase of the ARF family. This opens up the possibility to study the involvement of TTN5 in nutritional cell biology in the endomembrane system.

**Highlights:**

- TTN5 interacts with the large intracellular loop and variable region of IRON-REGULATED TRANSPORTER 1 (IRT1)
- TTN5 has a positive effect on root iron (Fe) reductase activity.
- TTN5 and IRT1 colocalize at the plasma membrane and in the endomembrane system related to vesicle transport
- TTN5 can interact with peripheral membrane proteins of the IRT1 interactome, EHB1, PATL2 and SNX1 suggesting a coordinating role in IRT1 regulation

**One-sentence summary:** TTN5, a small ARF-like GTPase, is connected to the dynamic regulation of IRON-REGULATED TRANSPORTER 1 (IRT1) in the vesicular trafficking system through direct protein interaction and colocalization, linking with various peripheral membrane proteins of the IRT1 interactome and iron reductase activity.

## Introduction

Plants adjust to environmental changes, such as variations of soil nutrient availability. Cellular membranes are very dynamic in plants, and this is crucial for sessile organisms. Plants can scan their environment for nutrients and control the activities of plasma membrane transporter proteins by vesicular trafficking. Many open questions still exist with regard to the environmental responsiveness and the complexity of the plant cell endomembrane system.

One essential element, iron (Fe), is abundant in the soil. But since Fe is mostly present in its ferric Fe^3+^ form in soil, it tends to be immobilized into insoluble complexes. Fe^3+^ must therefore be mobilized into a bioavailable form. Plants like *Arabidopsis thaliana* (Arabidopsis) are capable of this. One mechanism consists in reducing Fe^3+^ *via* plasma membrane FERRIC REDUCTION OXIDASE 2 (FRO2) (Robinson et al. 1999). The reactive ferrous Fe^2+^ is taken up by IRON-REGULATED TRANSPORTER 1 (IRT1) into root epidermis cells (Eide et al. 1996, Vert et al. 2002).

IRT1 is one of the founding members of an evolutionarily conserved divalent metal ion transporter family named ZIP (ZINC-REGULATED TRANSPORTER, ZRT, IRT-LIKE PROTEIN) family (Eide et al. 1996, Guerinot 2000). Divalent metal ions are potentially cytotoxic. The activity of IRT1, like that of other ZIP proteins, is controlled at the level of protein abundance at the plasma membrane (Connolly et al. 2002, Barberon et al. 2011, Shin et al. 2013, Barberon et al. 2014, Hu 2021). Importantly, IRT1 protein activity is likely controlled through transport from the *trans*-Golgi network/early endosomes (TGN/EE) to the plasma membrane, degradation by the lytic pathway or recycling back to the plasma membrane (Ivanov et al. 2014). However, the molecular machinery for this process is barely known.

The large cytosolic variable region (vr) between transmembrane domains three and four is a characteristic regulatory feature of ZIP proteins (Guerinot 2000, Gaither and Eide 2001). In IRT1, this regulatory variable region, IRT1vr, is crucial for the control of IRT1 abundance. IRT1vr has metal ion binding sites and residues that can be phosphorylated and ubiquitinated, which are processes that precede vesicular trafficking (Grossoehme et al. 2006, Kerkeb et al. 2008, Dubeaux et al. 2018, Cointry and Vert 2019). In a search for new components binding with IRT1vr we identified peripheral membrane proteins that directly or indirectly control the activity of IRT1 (Khan et al. 2019; Hornbergs et al. 2023). Among them is C2-domain protein ENHANCED BENDING 1 (EHB1) (Khan et al. 2019) which belongs to the ten-member calcium-dependent C2-DOMAIN ABSCISIC ACID-RELATED (CAR) protein family (EHB1 is also known as CAR6) (Rodriguez et al. 2014). CAR proteins act as tethers between the membrane and signaling or transport proteins that are involved in drought, defense or nutrition (Cui and Bauer 2023). They can also oligomerize and cause membrane deformations, making them candidate proteins involved in vesicle budding (Chen et al. 2023). EHB1 interacts with IRT1vr and it has an inhibitory effect on Fe acquisition (Khan et al. 2019). Another IRT1vr interactor is PATELLIN 2 (PATL2). It has a SEC14 domain coupled with a GOLD domain. The SEC14 domain forms a lipid-binding pocket that serves to present or transfer lipophilic substances at membranes (Montag et al. 2023). For example, PATL2 can bind α-tocopherol and it has been proposed that it prevents oxidative stress upon Fe acquisition, possibly by transferring and exchanging phospholipids and α-tocopherol at the membrane (Hornbergs et al. 2023). PATL2 may also initiate vesicle formation, as it binds proteins of the endomembrane system (Hornbergs et al. 2023). Another hint for the involvement of endomembrane trafficking in IRT1 protein control stems from the observation that SORTING NEXIN 1 (SNX1), a crucial component of the retromer complex for transmembrane protein sorting, binds to endomembrane structures containing IRT1, and it is a positive regulator of IRT1 recycling at the TGN (Ivanov et al. 2014). However, IRT1 does not seem to bind SNX1 directly.

Altogether, it looks like IRT1vr has a regulatory role as protein interaction platform to recruit components for vesicular trafficking of IRT1. Fe acquisition is affected by regulation of the vesicular transport mechanism, but how this is achieved and which proteins are involved are open questions that still remain largely elusive. Whether and how the various IRT1vr-interacting peripheral membrane proteins may be connected with each other to control localization of a plasma membrane protein like IRT1, is not clear yet.

Cycling of plasma membrane proteins is an important function of endomembrane trafficking (Valencia et al. 2016, Ivanov and Vert 2021). One protein family involved in this process in mammalian cells and yeast is the Ras superfamily of small GTPases. Small GTPases are important in vesicle-related processes as they function as molecular switches due to their GTPase activities to transmit signals. Small GTPases have usually low intrinsic nucleotide exchange and hydrolysis activity, and therefore, require the support of guanine nucleotide exchange factors (GEFs) and GTPase-activating proteins (GAPs). GEFs recruit the inactive, GDP-bound, GTPase to their site of action and lead to nucleotide exchange by GDP release. GTP binding leads to a conformational change of two regions referred to as switch I and II. The active, GTP-loaded, GTPases then exert their function, e.g. the recruiting and subsequent assembly of coat proteins by ARF1 or SAR1, until hydrolysis of the GTP by GAPs. Most of the known interactions occur in the active conformation of the GTPases (Sztul et al. 2019, Nielsen 2020, Adarska et al. 2021). Although the ARF-GTPase family is well-studied in mammals, there are only few well-established examples of an ARF-GTPase functioning in plasma membrane protein trafficking in plants, like that of ARF1-GNOM-mediated recycling of auxin carrier PIN FORMED 1 (PIN1), with GNOM being an ARF-GEF (Steinmann et al. 1999, Geldner et al. 2003). The presence of 21 ARF and ARF-like (ARL) GTPases encoded by the Arabidopsis genome raises the question whether other ARF members may act in endomembrane trafficking and what are their functions in the diverse physiological and nutrient assimilation pathways besides auxin signaling and cell division.

TITAN 5 (TTN5), also known as HALLIMASCH (HAL)/ARL2/ARLC1, is an ARF-like protein. It was identified in two independent screens for abnormal-embryo mutants arrested soon after egg cell division, indicating a central cellular function (Mayer et al. 1999, McElver et al. 2000), in agreement with its ubiquitous expression in plants (Mohr et al. 2024). TTN5 is more closely related in sequence to human ADP-ribosylation factor-like 2 (HsARL2) than any Arabidopsis protein. HsARL2 is associated with very diverse roles in cells, ranging from microtubule development, also identified for yeast and *Caenorhabditis* homologs (Bhamidipati et al. 2000, Fleming et al. 2000, Radcliffe et al. 2000, Antoshechkin and Han 2002, Tzafrir et al. 2002, Mori and Toda 2013), adenine nucleotide transport in mitochondria (Sharer et al. 2002) and control of phosphodiesterase activity in cilia (Ismail et al. 2011, Fansa and Wittinghofer 2016). HsARL2 is not only involved in a diverse set of functions and signaling cascades but it also binds very different protein partners for that. The same can be expected for TTN5. *TTN5* is expressed in the root epidermis (Mohr et al. 2024), like *IRT1* (Vert et al. 2002). TTN5 is present at the plasma membrane and in the endomembrane compartment. TTN5 is among the rare plant small GTPases for which GTPase activities are known. TTN5 can be considered atypical as it does not require a GEF to be in its active GTP-bound form and has a slow GTPase activity relying on a GAP (Mohr et al. 2024). Its unusually high intrinsic GDP-GTP exchange activity, that causes TTN5 to be present predominantly in a GTP-binding form, seems a prerequisite for its dynamic cellular localization and association with the dynamic endomembrane system and potential vesicular trafficking (Mohr et al. 2024). Physiological functions of this small GTPase, on the other hand, relevant during the entire plant life cycle, remain elusive until today.

A yeast two-hybrid screen had identified EHB1/CAR6 and PATL2 as IRT1 regulators (Khan et al. 2019, Hornbergs et al. 2023). We report here, that with this same strategy, we retrieved the HsARL2-related, small GTPase TTN5. We validated protein interactions of TTN5 and IRT1vr as well as the IRT1vr interactome and found a physiological Fe reductase phenotype linked with root Fe acquisition. Colocalization analysis indicated that active GTP-bound TTN5 was more colocalizing with IRT1 in intracellular vesicles than a mutant TTN5 form. Hence, our work indicates that the dynamic small GTPase TTN5 acts in a nutritional-environmental context in the vesicular trafficking system that controls the root Fe transporter IRT1.

## Results

### TTN5 interacts with the variable region of IRT1

To identify new cytoplasmic proteins regulating IRT1 we had developed a strategy to identify candidate interaction partners of IRT1vr (residues 145-192) and used IRT1vr as bait against a cDNA expression library prepared from Fe-deficient Arabidopsis roots (Khan et al. 2019, Hornbergs et al. 2023). We report here that 25 of the colonies (representing 25%) obtained in the reported yeast two-hybrid (Y2H) screen carried a fragment of the coding sequence of the gene AT2G18390, encoding the ARL-type small GTPase TTN5. The interaction was confirmed in a targeted Y2H assay (Figure 1A). Small GTPases can act as molecular switches in cells due to their rapid reactions and interactions with effectors, whereby TTN5 may not require a GEF for nucleotide exchange as it has high affinity for GTP (illustrated in Figure 1B) (Mohr et al. 2024). We further verified the protein interaction in plant cells using Bimolecular Fluorescence Complementation (BiFC) as a reconstitution of YFP in positively transformed cells that expressed the control marker mRFP (Figure 1C). We used previously characterized mutants of conserved amino acids in the GTP-binding pocket, Thr30-to-Asn (T30N, previously found to be dominant-negative or fast cycling) and Gln70-to-Leu (Q70L, with reduced GTP hydrolysis activity) (Figure 1B) (Mohr et al. 2024). Both, the cYFP-TTN5^T30N^ and cYFP-TTN5^Q70L^ forms interacted with nYFP-IRT1vr (Figure 1D-E). The BiFC interactions were specific as neither protein partner was able to bind negative controls in this assay (Figure 1F-G). We further elucidated whether IRT1vr might discriminate between the conserved TTN5 variant forms using a quantitative analysis of the interaction by Förster Resonance Energy Transfer-Acceptor Photobleaching (FRET-APB). This approach is based on close-proximity-dependent energy transfer of an excited GFP-tagged TTN5 donor to a mCherry acceptor, here IRT1vr-mCherry. Energy transfer was quantified as FRET efficiency (Figure 1H-J). FRET efficiencies between all GFP-TTN5 forms and IRT1vr-mCherry were significantly higher compared to the donor-only sample, confirming protein interactions between all TTN5 forms and IRT1vr. As a final evidence, we performed an *in planta* pulldown experiment to show interaction of full-length IRT1 and TTN5 (Figure 1K) using an Arabidopsis plant line expressing IRT1-mCherry together with hemagglutine-tagged (HA)-TTN5. The mCherry part was integrated in the first small cytosolic loop of IRT1 at position 80 (proIRT1::IRT1Q80-mCherry, briefly IRT1-mCherry), and this fusion protein was functional as it fully complemented the severe *irt1-1* (SALK_054554) Fe deficiency phenotype (Fukao et al. 2011) (Supplemental Figure S1). HA_3_-TTN5 (pro35S::HA_3_-TTN5, briefly HA_3_-TTN5) was functional as it complemented the *ttn5-1* phenotype as previously described (Mohr et al. 2024). We pulled down protein complexes from whole seedlings using anti-HA beads. This resulted in an increased protein abundance of HA_3_-TTN5 (28 kDa) in the elution fraction, as expected (Figure 1K). Importantly, IRT1-mCherry was simultaneously coprecipitated and detected in this elution fraction while it was not detected in a control pulldown experiment without HA_3_-TTN5 (Figure 1K), indicating that the two full-length proteins interact closely in Arabidopsis plant cells.

**Figure 1.**
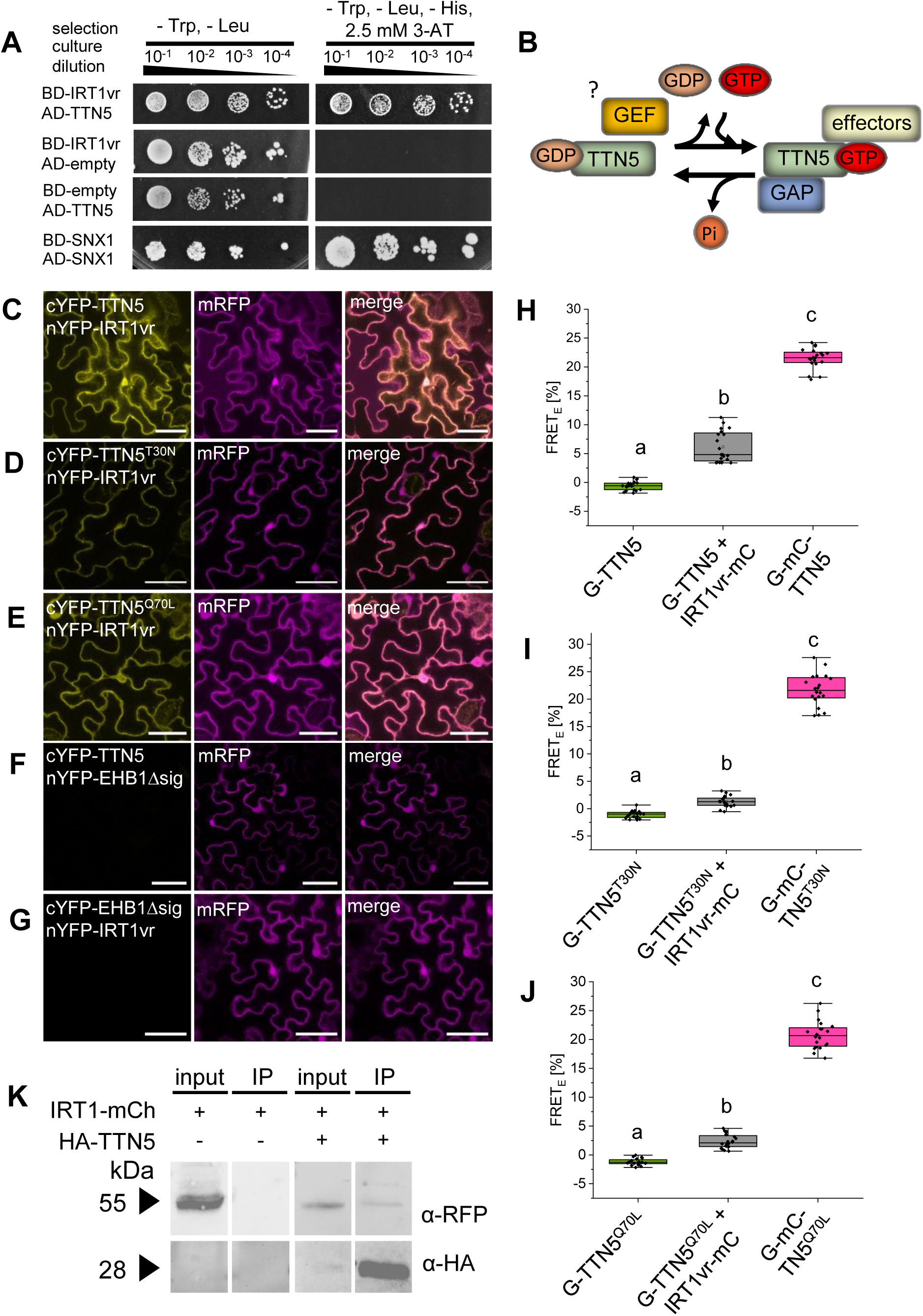
TTN5 interacts with IRT1 via the cytosolic loop and variable region IRT1vr. (A), Targeted Y2H assay between BD-IRT1vr (IRT1-variable region, vr) and AD-TTN5. Yeast cotransformation samples were spotted in 10-fold dilution series (A_600_ = 10^1^-10^4^) on double selective (LW; control) and triple selective +0.5 mM 3-AT (LWH, selection for protein interactions) SD-plates. SNX1/SNX1, positive control (Pourcher et al. 2010), BD-IRT1vr/AD-empty and BD-empty/AD-TTN5 respective negative controls. (B), Schematic representation of the switching mechanism of small GTPases based on TTN5. TTN5 can switch between an inactive GDP-loaded form to an active GTP-loaded one. Many small GTPases require a guanidine exchange factor (GEF) for nucleotide exchange (Vernoud et al. 2003). However, TTN5 has unusually high intrinsic GDP to GTP nucleotide exchange capability and help by a GEF may not be needed (indicated by space between TTN5 and GEF). Nevertheless, it is not excluded that TTN5 can interact with a GEF (indicated by a question mark) in plant cells. GTP hydrolysis has to be catalyzed by a GTPase-activating protein (GAP) (Mohr et al. 2024). GTP-loaded TTN5 can interact with effector proteins for signal transition. (C-G), Validation of TTN5-IRT1vr interaction by bimolecular fluorescence complementation (BiFC) of split YFP. Proteins of interest are fused either to the C- or N-terminal half of split-YFP in this assay. Potential interaction is indicated by a reconstituted YFP signal. mRFP serves as a transformation control. (C-E), nYFP-IRT1vr showed YFP complementation in combination with (C) cYFP-TTN5, (D) cYFP-TTN5^T30N^ and cYFP-TTN5^Q70L^ variants. The complementation takes place in nucleus and cytoplasm predominantly. (F-G), The combinations (F) cYFP-TTN5/nYFP-EHB1Δsig and (G) cYFP-EHB1Δsig/nYFP-IRT1vr served as negative controls. Each combination was tested a minimum of three times with comparable results. Scale bar 50 µm. (H-J), Confirmed interaction via Förster resonance energy transfer (FRET)-acceptor photobleaching (APB). Significant increase of FRET-efficiency (FRET_E_) between IRT1vr-mCherry (IRT1vr-mC) and (H) GFP-TTN5 (G-TTN5), (I) GFP-TTN5^T30N^ (G-TTN5^T30N^) or (J) GFP-TTN5^Q70L^ (G-TTN5^Q70L^) indicate short protein distances. GFP-fusions are donor-only samples and serve as a negative control. GFP-mCherry-tagged constructs show intra-molecular FRET as a respective positive control. Each combination was tested a minimum of three times with comparable results with a minimum of ten individual measurements (n ≥ 10). The box represents the 25–75^th^ percentiles, and the median is indicated. The whiskers show the 5^th^ and 95^th^ percentiles. One-way ANOVA with Tukey post-hoc test was performed. Different letters indicate statistical significance (p < 0.05). (K) *In planta* coimmunoprecipitation (Co-IP) experiment demonstrating interaction between full-length IRT1-mCherry (proIRT1::IRT1Q80-mCherry) and HA_3_-TTN5 (pro35S::HA_3_-TTN5) using Arabidopsis roots carrying both transgenes or only IRT1-mCherry grown in parallel under Fe-deficient conditions. Following protein complex extraction, coimmunoprecipitation was conducted with anti-HA antibody. HA_3_-TTN5 protein was enriched by the pull-down, resulting in an increased band intensity in the elution (IP) fraction compared to the input. IRT1Q80-mCherry protein was copurified with HA_3_-TTN5 in two out of three repetitions. No coimmunoprecipitation is detected in the negative control in the absence of HA_3_-TTN5 protein.

In sum, we can conclude that TTN5 is an interactor of IRT1 with IRT1vr as interaction surface. We hypothesize that TTN5 is a potential link in the coordination of IRT1 regulation with vesicle trafficking typical for ARF GTPases.

### TTN5 affects Fe homeostasis

Following up on the IRT1-TTN5 interaction, we tested whether TTN5 might affect plant growth under Fe-deficient conditions and/or the regulation of the Fe deficiency response. Homozygous *ttn5* knock-outs are embryo-lethal (Mayer et al. 1999, McElver et al. 2000), and initial growth experiments with heterozygous plants (*ttn5-1^+/-^*) showed no obvious developmental defects in comparison with wild type sibling plants under regular growth conditions (Figure 2A, Supplemental Figure S2; (Mohr et al. 2024)). Likewise, when plants were germinated in Fe-sufficient and -deficient conditions, we did not note any difference in root length. Both, wild type and *ttn5-1^+/-^* siblings, had increased root length under Fe-deficient conditions (Figure 2B), as expected in our growth system (Gratz et al. 2019), but there was no difference between them. SPAD values indicating chlorophyll contents were also similar in the wild type and *ttn5-1^+/-^*seedlings, indicating that the *ttn5* mutation has no effect on leaf color (Figure 2C). Also at advanced growth stages, *ttn5-1^+/-^* plants did not differ in growth and leaf color from their wild type siblings, neither in in control soil (pH 6.2) nor in an alkaline, calcareous soil (ACS, pH 8). The latter conditions require Fe mobilization capacities since Fe acquisition is inhibited at high pH values in the presence of bicarbonates, resulting in smaller plants with leaf chlorosis ((Ohwaki and Sugahara 1997, Schmid et al. 2014), Supplemental Figure S2B). Both wild type and *ttn5-1^+/-^*plants showed similarly lower aerial biomass increase and lower chlorophyll on ACS conditions over a period of 43 days compared to the control conditions (Supplemental Figure S2A-D). We examined the heterozygous seedlings at the molecular and physiological level. *TTN5* expression was nearly 50% reduced in *ttn5-1^+/-^* compared to wild type siblings, consistent with single allele presence (Figure 2D). Very interestingly, we detected a 15% decrease of *TTN5* gene expression under Fe deficiency compared with Fe sufficiency in both, wild type and *ttn5-1^+/-^*, indicating that TTN5 might be perhaps less needed under low Fe supply (Figure 2D). *FRO2* encodes an Fe uptake component, the ferric reductase oxidase enzyme (Robinson et al. 1999), that closely interacts with IRT1 in the root plasma membrane (Martín-Barranco et al. 2020). *FRO2* and *IRT1* gene expression increase in response to Fe deficiency, and this serves as a molecular marker for the root Fe deficiency status, e.g. (Gratz et al. 2019). No difference of gene expression levels of *FRO2* and *IRT1* occurred in *ttn5-1^+/-^* roots compared with wild type, and both genes were induced under low Fe versus high Fe as expected (Figure 2E-F). This indicated that root Fe deficiency responses might not be compromised in *ttn5-1^+/-^*. However, we then noted that in two out of three experiments (each including the indicated biological replicates), root Fe reductase activity was altered. It was up-regulated under Fe-deficient versus-sufficient conditions in wild type and *ttn5-1^+/-^* roots, as expected, e.g. (Robinson et al. 1999, Gratz et al. 2019) but the induction was lower in *ttn5-1^+/-^* compared with wild type siblings (Figure 2G).

**Figure 2.**
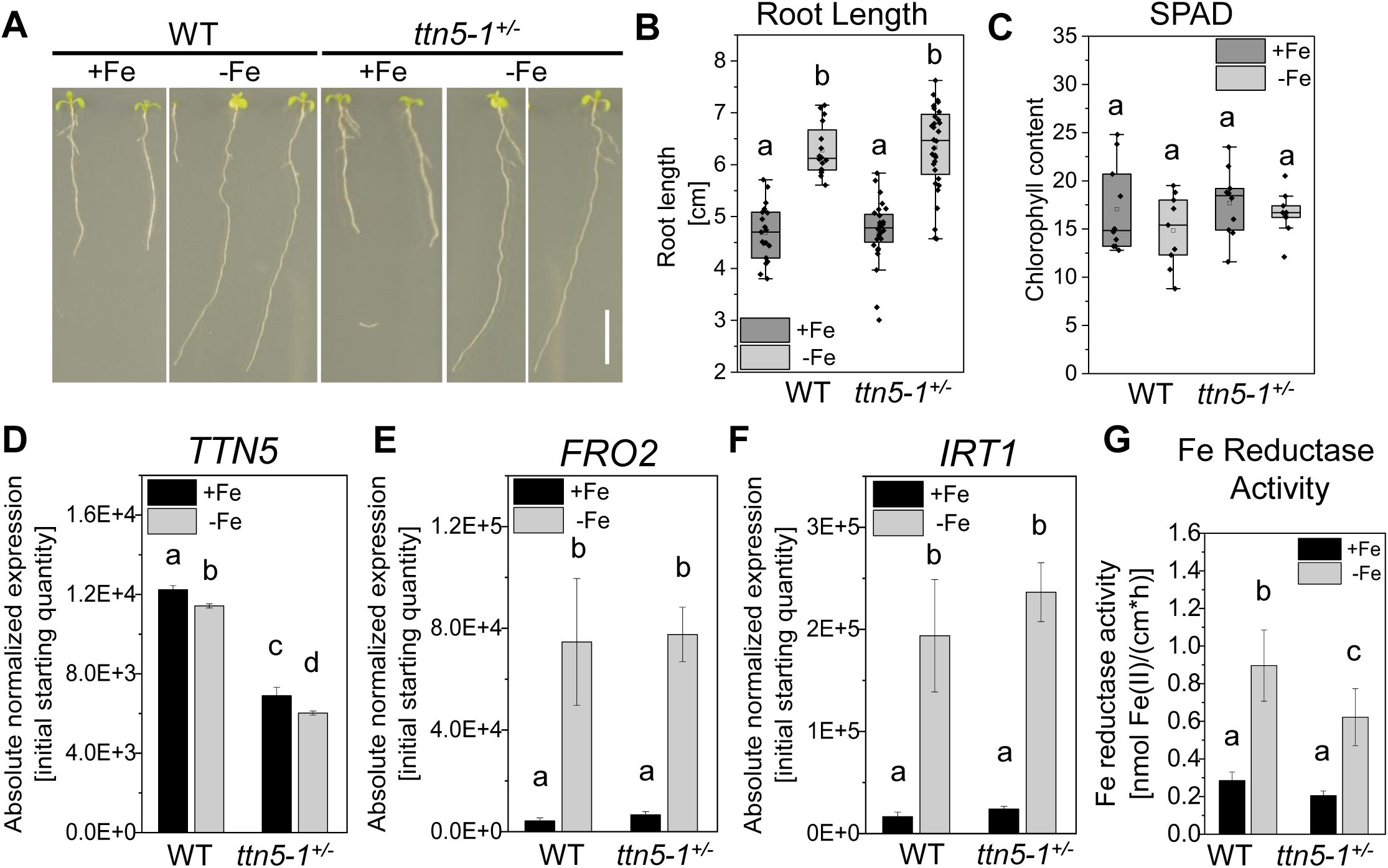
Heterozygous *ttn5-1^+/-^* seedlings can have an Fe reductase phenotype. Heterozygous *ttn5-1^+/-^*and their wild type sibling seedlings were grown on +/-Fe Hoagland plates for ten days (A, B) or in the two-week system (C-H). All plants were individually genotyped as wild type or *ttn5-1^+/-^*. Note that homozygous *ttn5-1^+/+^*are embryo-lethal. (A), + and -Fe-grown *ttn5-1^+/-^* seedlings appeared phenotypically as wild type (WT) siblings. Scale bar 1 cm. (B), Quantified root length of (A). Root length increased under Fe-deficient conditions (-Fe, light grey boxes) compared to sufficient conditions (+Fe, dark grey boxes), similar in *ttn5-1^+/-^* and wild type siblings (+Fe: WT n = 19; *ttn5-1^+/-^* n = 28; -Fe: WT n = 16; *ttn5-1^+/-^*n = 32). The box represents the 25–75^th^ percentiles, and the median is indicated. The whiskers show the 5^th^ and 95^th^ percentiles. (C), Soil Plant Analysis Development (SPAD) values, reflecting chlorophyll amounts, were similar under both Fe conditions (-Fe, light grey boxes; +Fe, dark grey boxes) and comparable between the plants (+Fe: n = 10; -Fe: n = 9). The box represents the 25–75^th^ percentiles, and the median is indicated. The whiskers show the 5^th^ and 95^th^ percentiles. (D-F), Gene expression data determined by RT-qPCR of *TTN5, FRO2* and *IRT1* in *ttn5-1^+/-^* roots compared to wild type siblings in Fe-sufficient (+Fe, black bars) or -deficient (-Fe, grey bars) conditions. (D), *TTN5* expression levels were higher in +Fe than -Fe conditions in wild type and *ttn5-1^+/-^* seedlings. Expression was about half in mutant versus wild type (E-F) *FRO2* and *IRT1* expression was induced upon Fe deficiency versus sufficiency in wild type and *ttn5-1^+/-^* seedlings in comparable amounts. Data was obtained from three biological replicates (n = 3). (G), Root ferric reductase activity determined by the Fe^2+^-ferrozine assay at 562 nm. Iron reductase activity was increased upon Fe deficiency in wild type. Same was found in *ttn5-1^+/-^* sibling seedlings under Fe deficiency but to lesser extent. The assay was performed with three (wild type) and four (*ttn5-1^+/-^*) biological replicates (WT n = 3; *ttn5-1^+/-^* n = 4). The experiment was repeated in total three times. In two cases, Fe reductase activity was reduced in *ttn5-1^+/-^*compared with wild type siblings (one representative experiment shown), and once it was not different. One-way ANOVA with Tukey as post-hoc test was performed. Different letters indicate statistical significance (p < 0.05).

Taken together, heterozygous *ttn5-1^+/-^* seedlings have an Fe reductase phenotype that indicates a positive effect of TTN5 at the physiological level on Fe acquisition responses in the plasma membrane. This was interesting, because the two other interactors of IRT1vr, EHB1/CAR6 and PATL2, had an opposite effect on Fe reductase activity (Khan et al. 2019, Hornbergs et al. 2023).

### TTN5 colocalizes with IRT1 in the plasma membrane and in vesicles

To investigate whether TTN5 may be located in similar places as IRT1 in root cells, as expected for interacting proteins, we grew and compared side by side localization of fluorescent signals in Arabidopsis plant roots expressing either IRT1-mCitrine (proIRT1::IRT1-mCitrine/*irt1-1*), described in (Dubeaux et al. 2018), or YFP-TTN5 (pro35S::YFP-TTN5), described in (Mohr et al. 2024). A similar approach has been used to compare intracellular localization of IRT1 and PATL2 fusion proteins in root cells (Hornbergs et al. 2023). As previously described, fluorescent signal of IRT1-mCitrine was present at the plasma membrane of root epidermis cells in our growth system (Supplemental Figure S3A) (Dubeaux et al. 2018, Hornbergs et al. 2023). Fluorescent plasma membrane signal was also detected for YFP-TTN5 expressing seedlings (Supplemental Figure S3B), as corresponding to the previously identified pattern (Mohr et al. 2024). Virtual cross sections of both roots highlighted the described polar localization for IRT1-mCitrine, visible in only a very few epidermis cells, as previously seen (Supplemental Figure S3C; (Dubeaux et al. 2018, Hornbergs et al. 2023) compared with an equally distributed YFP localization in the plasma membrane of YFP-TTN5 expressing seedlings in the epidermis (Supplemental Figure S3D). Next to plasma membrane localization both, fluorescent signals of IRT1-mCitrine and YFP-TTN5 expressing seedlings, were also present in vesicle-like structures in root cells (Supplemental Figure S3E-F). These findings suggest that indeed, IRT1 and TTN5 may locate to similar structures inside root cells, supporting an interaction at these places.

To confirm the obtained Arabidopsis data and to investigate potential colocalization of TTN5 with IRT1 in the same cells, we used the transient expression system of *Nicotiana benthamiana* (*N. benthamiana*). This system has the advantage that fusion proteins of TTN5 and IRT1 are well detectable, and through well-established pharmacological approaches the localization patterns of TTN5 and IRT1 proteins can be confirmed (Ivanov et al. 2014, Mohr et al. 2024). We used a GFP-tagged IRT1 with GFP inserted inside IRT1vr (hereafter termed IRT1-GFP). This IRT1 fusion protein was present at the plasma membrane. Fluorescent spots were also found in close proximity to the plasma membrane (Supplemental Figure S4A), consistent with previous reports (Barberon et al. 2011, Ivanov et al. 2014). To further evaluate its usefulness for localization studies, we compared IRT1-GFP signals with already described C-terminally-tagged IRT1-mCherry signals in *N. benthamiana* pavement cells before and after wortmannin treatment, see (Ivanov et al. 2014). Wortmannin is a fungal metabolite with an inhibitory effect on phosphatidylinositol-3-kinase (PI3K) function, leading to swollen multivesicular bodies (MVBs) and therefore is commonly used to investigate endocytosis (Cui et al. 2016). Signals upon expression of these two fusion proteins showed colocalization (Supplemental Figure S4B) and, following wortmannin treatment, were observed among others in swollen structures, suggesting partial presence in the MVBs (Supplemental Figure S4C), indicating that the IRT1-GFP fusion protein was viable for IRT1 localization studies. IRT1-GFP also partially complemented an early rosette growth deficit of *irt1-1* (in pro35S::IRT1-GFP/*irt1-1* plants), indicating partial functionality of the IRT1-GFP construct for this trait (Supplemental Figure S4D).

We detected colocalization of IRT1-GFP and mCherry-TTN5 including TTN5-mutant fluorescent signals at the plasma membrane or just below (Figure 3A-C). Previously, we had described YFP-TTN5 localization in vesicle-like structures and close proximity of YFP-TTN5 to the plasma membrane (Mohr et al. 2024). Moving fluorescent structures upon YFP-TTN5^T30N^ expression were less mobile than for YFP-TTN5 and YFP-TTN5^Q70L^ (Mohr et al. 2024). Very interestingly, the different mCherry-TTN5 forms have different abilities to colocalize with IRT1 by Pearson coefficients (Figure 3D). While Pearson coefficients of signals of IRT1-GFP with mCherry-TTN5 and mCherry-TTN5^Q70L^ were 0.80 and 0.77, it was higher, namely 0.90, for IRT1-GFP and mCherry-TTN5^T30N^, demonstrating higher general colocalization between IRT1-GFP and the low-mobile TTN5^T30N^ form (similar results for overlap coefficients Supplemental Figure S4E). Signals of all three mCherry-TTN5 forms were also localized in IRT1-GFP-positive structures close to the plasma membrane, suggesting colocalization in plasma membrane-derived vesicles (Figure 3E-G). Surprisingly, the Pearson coefficients changed when these spotted structures were analyzed (Figure 3H; Supplemental Figure S4F similar results for overlap coefficients). While the values for fluorescent signals of mCherry-TTN5 (0.80) and mCherry-TTN5^Q70L^ (0.78) constructs with IRT1-GFP remained relatively the same, we obtained with 0.78 now a similar coefficient with mCherry-TTN5^T30N^. Similar results were achieved while checking the overlaps of mCherry-TTN5 structures with IRT1-GFP and vice versa with a centroid-based analysis. We obtained overlaps of 36-41% for mCherry-TTN5 signals with IRT1-GFP (Supplemental Figure S4G). 54% of IRT1-GFP-positive structures colocalized with signals of mCherry-TTN5^T30N^, whereas 64% and up to 83% colocalized with mCherry-TTN5 and mCherry-TTN5^Q70L^, respectively. From this, we hypothesized that IRT1 is more likely to be colocalized with TTN5^T30N^ at the plasma membrane than with TTN5 wild type or TTN5^Q70L^. as the latter may be active in promoting intracellular cycling. This was consistent with the previous observation that signals of mCherry-TTN5^T30N^ were moving slower in certain cells than those of the other two TTN5 fusion proteins. We therefore deduce that TTN5^T30N^ may slow down the removal of IRT1 from the plasma membrane. In an attempt to identify the intracellular structures in which colocalization of TTN5 and IRT1 occurred, we treated the cells with wortmannin (Figure 3I). Structures positive for both IRT1-GFP and mCherry-TTN5 fluorescent signals became doughnut-shaped suggesting IRT1-TTN5 colocalization in MVBs (Figure 3J-K). Presence of IRT1 in these late endosomal compartments has been observed previously, and might be part of the pathway to IRT1 vacuolar degradation (Dubeaux et al. 2018), same for mCherry-TTN5 (Mohr et al. 2024). Thus, our findings emphasize the involvement of TTN5 in the endomembrane trafficking of the Fe transporter IRT1.

**Figure 3.**
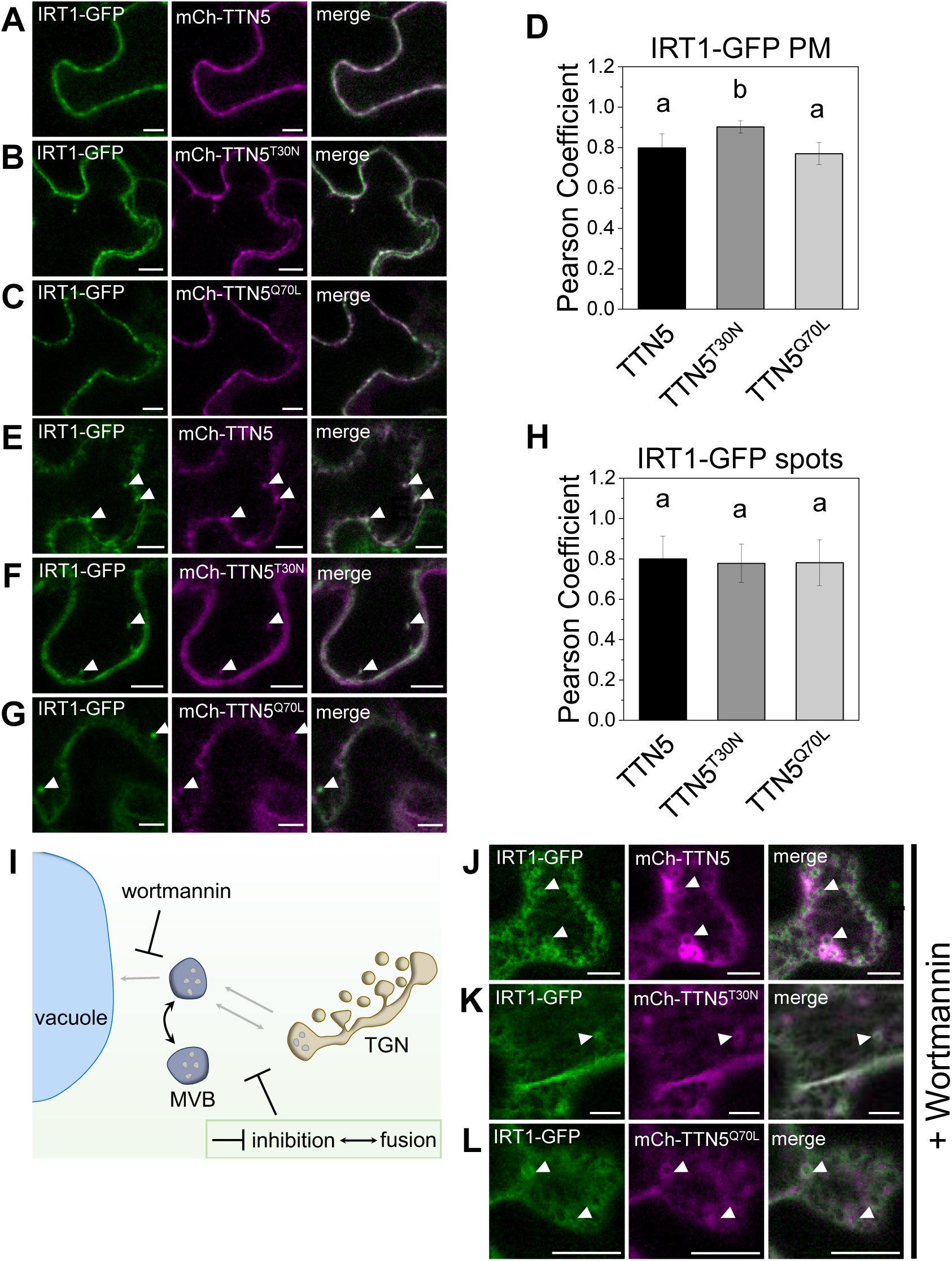
TTN5 and IRT1 fluorescence protein signals colocalize at the plasma membrane and in multi-vesicular bodies (MVBs) in *N. benthamiana* leaf pavement cells. Fluorescent signals of mCherry-TTN5 and its mutant derivatives mCherry-TTN5^T30N^ and mCherry-TTN5^Q70L^ were colocalized with IRT1-GFP (pro35S::IRT1-GFP) expression in *N. benthamiana* leaf pavement cells. (A-C), Fluorescent signals of mCherry-TTN5 and mCherry-TTN5 variants expressing constructs were present at the plasma membrane and in distinct spots together with IRT1-GFP signal. (D), JACoP-based colocalization analysis of mCherry-TTN5 and variants with IRT1-GFP (Bolte and Cordelières, 2006) performed with ImageJ (Schneider et al. 2012). Comparison of Pearson’s coefficients for IRT1-GFP with mCherry-TTN5, mCherry-TTN5^T30N^ and mCherry-TTN5^Q70L^ at the plasma membrane (PM). Fluorescent signals of mCherry-TTN5^T30N^ construct were statistically significant more present at the PM compared to signals of mCherry-TTN5 and mCherry-TTN5^Q70L^. (E-G), Internalized fluorescent signals of IRT1-GFP expression were present in vesicle-like structures positively labelled by mCherry-TTN5 and mCherry-TTN5 variant signals. (H), Comparison of Pearson’s coefficients for IRT1-GFP with mCherry-TTN5, mCherry-TTN5^T30N^ and mCherry-TTN5^Q70L^ in internalized spots. Fluorescent signals of all mCherry-TTN5 construct colocalized in a similar way with IRT1-GFP signals. (I), Schematic illustration of the inhibiting function of the fungal metabolite wortmannin on cellular trafficking routes between the vacuole and the *trans*-Golgi network via MVBs, leading to swelling of MVBs. (J-L), Cells were treated with wortmannin (10 µM). Swollen MVBs are visible with the GFP- and mCherry-channel, respectively. Colocalization is indicated with filled arrowheads. Microscopic experiments were conducted in a minimum of three repetitions (n ≥ 3). JACoP analyses were performed three times (n = 3). One-way ANOVA with Fisher-LSD as post-hoc test was performed. Different letters indicate statistical significance (p < 0.05). Scale bar 10 µm.

### TTN5 interacts with IRT1 regulators EHB1, SNX1 and PATL2

IRT1 can associate and colocalize with peripheral membrane proteins SNX1, EHB1 and PATL2 that can be associated with events of vesicular trafficking (Ivanov et al. 2014, Khan et al. 2019, Hornbergs et al. 2023). This opened up the possibility that TTN5 may associate not only with IRT1 but also with proteins of IRT1 interactome in close proximity in cells, prompting us to investigate this further.

At first, we tested IRT1-interacting EHB1. CAR proteins like EHB1 have a plant-specific insertion of an extra region within the C2 domain known as signature (sig) domain, that is involved in binding proteins including IRT1 (Rodriguez et al. 2014, Khan et al. 2019). We demonstrated above that nYFP-EHB1Δsig did not interact with cYFP-TTN5 in BiFC (Figure 1F). However, when using for interaction with cYFP-TTN5 the intact nYFP-EHB1 protein or the EHB1 sig-nYFP domain alone we did find evidence of protein interaction in close proximity to the plasma membrane using BiFC in contrast to EHB1Δsig (Figure 4A-C). Surprisingly, the cYFP-TTN5 mutant forms both failed to complement with nYFP-EHB1 in the split-YFP assay (Figure 4D-E). This was surprising because FRET-APB experiments validated the interaction between EHB1-GFP as the donor and TTN5-mCherry as well as of the two TTN5 mutant variants as the acceptors (Figure 4F). Surprisingly, the FRET pairs EHB1-GFP with TTN5^T30N^-mCherry and TTN5^Q70L^-mCherry resulted in an even higher FRET-efficiency compared to TTN5-mCherry (Figure 4F). Perhaps, the mutant behavior and difference between BiFC and FRET-APB were due to the different tag orientations in these two types of assays.

**Figure 4.**
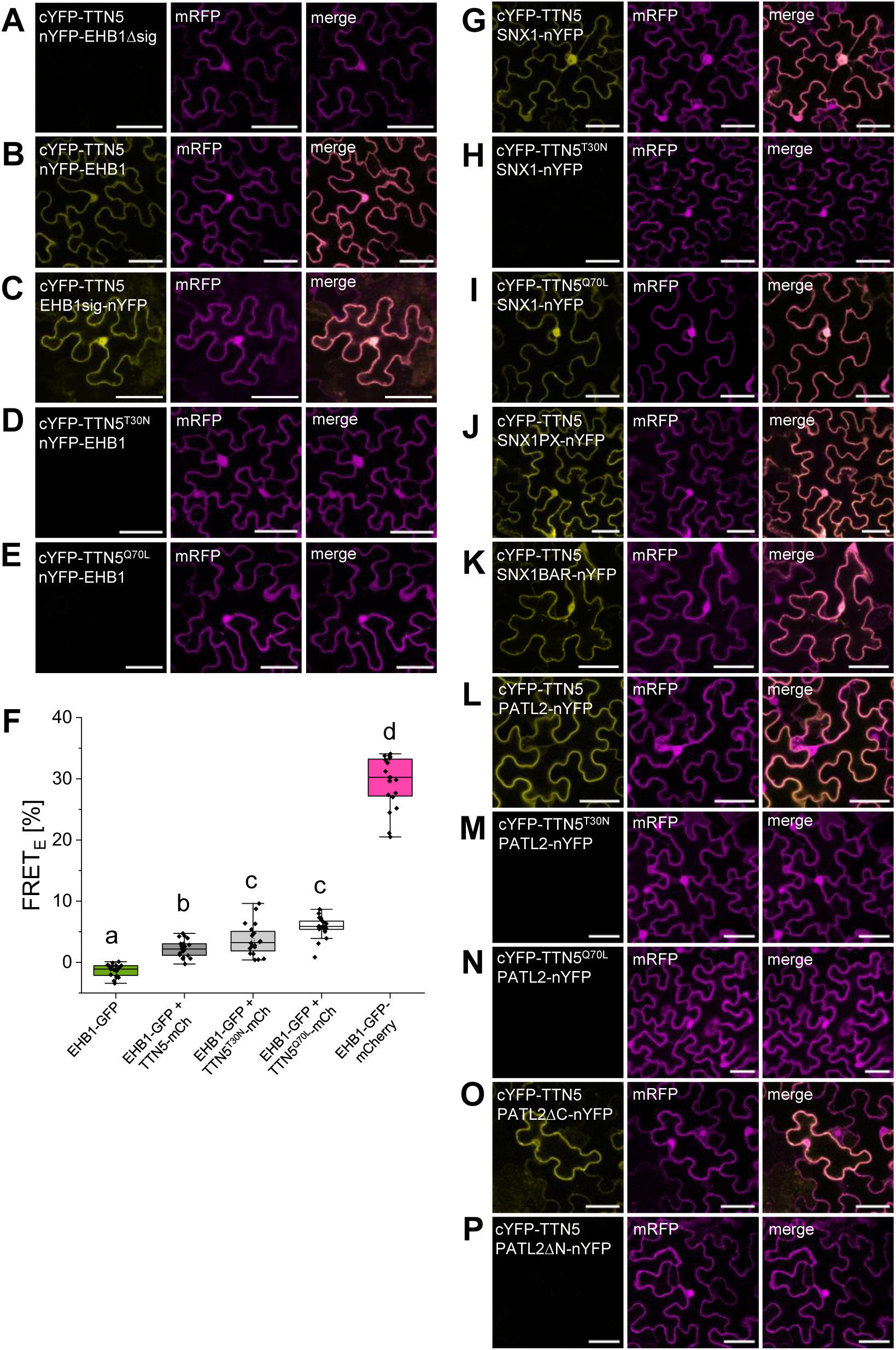
TTN5 interacts with IRT1 regulators EHB1, SNX1 and PATL2. (A-E, G-P), Bimolecular fluorescence complementation (BiFC) of split YFP experiments with TTN5 and known IRT1 regulators EHB1 (Khan et al. 2019), SNX1 (Ivanov et al. 2014) and PATL2 (Hornbergs et al. 2023). mRFP expression serves as a transformation control. (A-E), TTN5 interacts with EHB1 via the sig domain. (B) TTN5 and EHB1 showed complementation whereas (D, E) variants were not able to complement split-YFP. (A, C) TTN5 was tested together with EHB1sig (signature domain, sig domain only) and with EHB1 harboring a deletion of the CAR sig domain (sig domain defined in Rodriguez et al. (2014), Khan et al. (2019), EHB1Δsig). Deletion of the sig domain resulted in loss of complementation. (F), Interaction between EHB1-GFP (donor) with TTN5-mCherry and TTN5-mCherry variants (acceptor) was confirmed by FRET-APB. EHB1-GFP as donor-only sample (negative control), EHB1-GFP-mCherry for intra-molecular FRET (positive control). In this assay, TTN5 variants interacted with EHB1. Each combination was tested a minimum of three times with a minimum of ten individual measurements (n ≥ 10) with comparable results. The box represents the 25–75^th^ percentiles, and the median is indicated. The whiskers show the 5^th^ and 95^th^ percentiles. (G-K), BiFC showing that TTN5 interacts with SNX1. (G) TTN5 and (I) TTN5^Q70L^ showed complemented YFP signal together with SNX1. No YFP fluorescence was detectable for (H) TTN5^T30N^ with SNX1. (J, K), An interaction took place between TTN5 and the (J) PX and (K) the BAR domain of SNX1. (L-P), BiFC showing that TTN5 interacted with PATL2. (L) PATL2 showed complemented YFP signal with N-terminally-tagged TTN5. (M) TTN5^T30N^ and (N) TTN5^Q70L^ did not complement YFP with PATL2. (O-P), Deletion mutants of PATL2 showed dependency of the N-terminus for complementation with TTN5. (O) YFP signal was visible for PATL2 N-terminus (PATL2ΔC). (P) On the other hand, the C-terminus alone (PATL2ΔN) had no detectable YFP. Every construct was tested a minimum of three times (n ≥ 3) with similar results. Scale bar 50 µm. One-way ANOVA with Tukey post-hoc test was performed. Different letters indicate statistical significance (p < 0.05).

In another BiFC experiment, cYFP-TTN5 was able to complement split-YFP together with SNX1-nYFP close to the plasma membrane (Figure 4G). The cYFP-TTN5^T30N^ mutant failed in BiFC (Figure 4H), while the hydrolysis impaired TTN5 GTPase variant interacted with SNX1-nYFP (Figure 4I). SNX proteins are classified by their characteristic domains. SNX1 represents the class I SNX proteins, characterized by the presence of PHOX-homology (PX) and Bin-Amphiphysin-Rvs (BAR) domains (Heucken and Ivanov 2018). Such a domain organization is typical for proteins that function in membrane and phosphoinositide binding by the PX domain, whereas the BAR domain is involved in membrane curvature sensing and the formation of endosomal tubules (Peter et al. 2004, Frost et al. 2009). Interestingly, ARF-GAPs which are needed for the GTP hydrolysis, contain a BAR domain as well (Memon 2004). We found evidence that cYFP-TTN5 binds to both the SNX1 PX-nYFP and BAR-nYFP domains using BiFC (Figure 4J-K).

Lastly, plasma membrane-localized SEC14 protein PATL2 binds IRT1vr (Hornbergs et al. 2023). We tested interaction ability of cYFP-TTN5 with PATL2-nYFP in BiFC experiments. YFP signals indicating complementation and interaction were visible at the plasma membrane, fitting to the localization studies of both PATL2 and TTN5 (Figure 4L). We were not able to observe YFP signal complementation for the combination of PATL2-nYFP and any of the two GTPase variants cYFP-TTN5^T30N^ (Figure 4M) and cYFP-TTN5^Q70L^ (Figure 4N). This was very interesting as it is either a limitation of the method or could indicate that the TTN5 switching mechanism may be required for a potential interaction between the proteins. However, this needs to be tested further in the future. To further identify the region of interaction, we tested already published deletion mutants of PATL2 (Montag et al. 2020). PATL2 can be divided into two protein halves, consisting of either an N-terminal part with no particular conformation or the C-terminal part with CRAL-TRIO-N-terminal extension (CTN)-SEC14-Golgi dynamics (GOLD) domains that is conserved between SEC14-GOLD proteins in plants (Montag et al. 2020). The N-terminal part is needed for the interaction with IRT1vr (Montag et al. 2020). We used the PATL2 deletion mutants PATL2ΔC lacking the CTN-SEC14-GOLD part and PATL2ΔN lacking the IRT1vr-interacting N-terminal part. PATL2ΔC-nYFP together with cYFP-TTN5 complemented YFP fluorescence indicating interaction (Figure 4O), whereas no signal was obtained by PATL2ΔN-nYFP in combination with cYFP-TTN5 (Figure 4P), similar as in the case of IRT1vr (Hornbergs et al. 2023). Interestingly, the fluorescent complementation signal between cYFP-TTN5 and PATL2ΔC-nYFP was no longer present at the plasma membrane consistent with the observation that the PATL2 plasma membrane localization is dependent on C-terminal PATL2 domain region (Montag et al. 2020).

We also analyzed *EHB1, SNX1* and *PATL2* gene expression. All genes were expressed in similar levels in both wild type and *ttn5-1^+/-^* (Supplemental Figure S5A-C) indicating that there was no feedback at the level of regulation of genes encoding the TTN5 interactors.

In summary, we could show that TTN5 is not only able to interact with IRT1vr but also with three peripheral membrane proteins that can be present in the vicinity of IRT1 at membrane sites in root cells.

## Discussion

Here, we report that the small GTPase TTN5 is a potential link in IRT1-related Fe homeostasis and trafficking of the transporter within the endomembrane system (Figure 5). Such a finding is highly interesting. It was known that IRT1 is post-translationally controlled involving the vesicular trafficking system, but components that may bind with IRT1 during this process had remained obscure. Moreover, the connections of small GTPases like TTN5 with endomembrane trafficking were mostly not ascribed to concrete physiological roles. Hence, our report is very timely, and highlighting direct protein connections between cargo and vesicle trafficking components fills a gap in our understanding.

**Figure 5.**
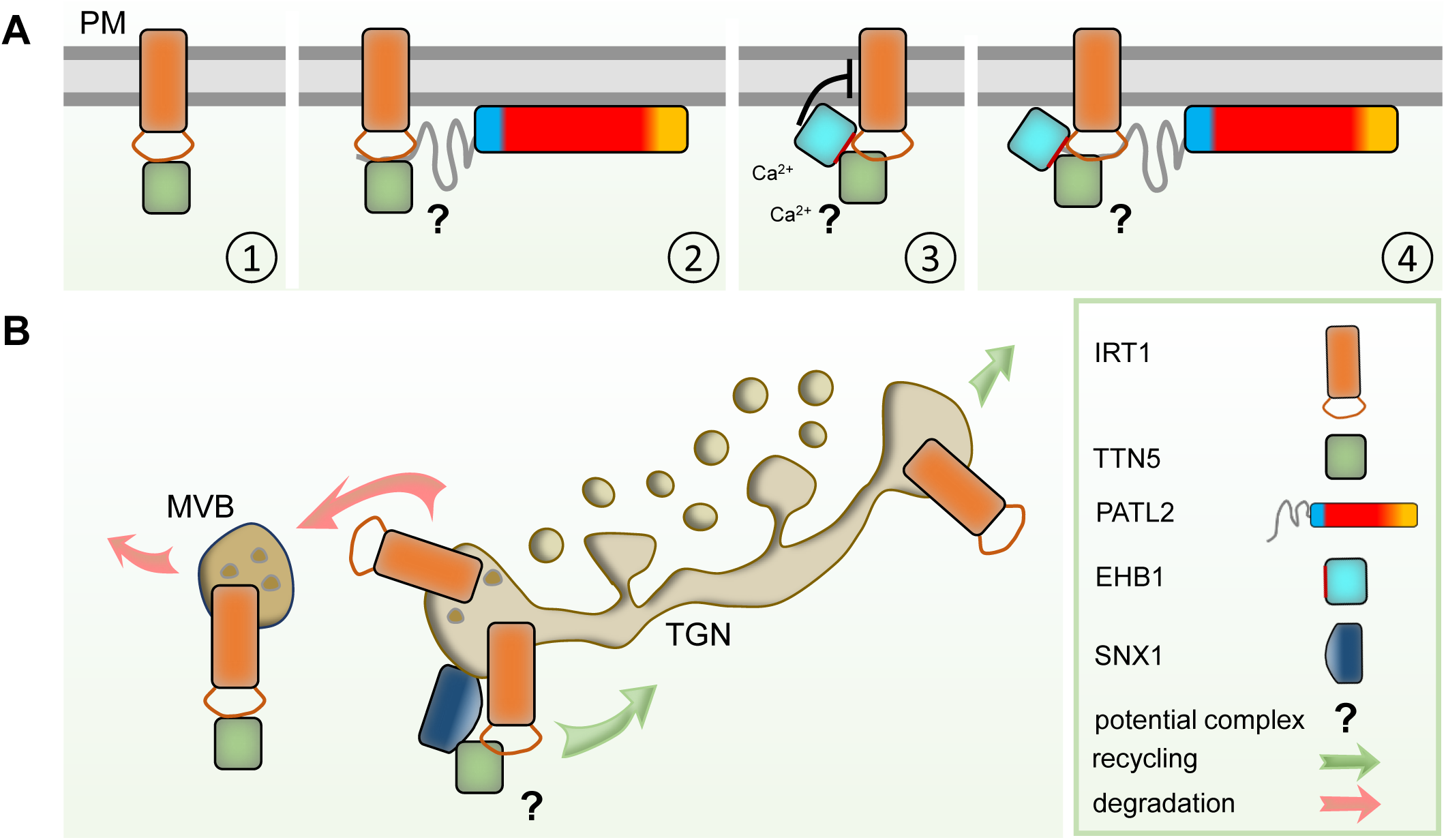
Summary model: Action of TTN5 within the IRT1-regulatory network. A schematic overview of the identified TTN5 interactors within the IRT1-regulatory network (A) at the plasma membrane (PM) or (B) in the endomembrane compartments of the *trans*-Golgi network (TGN) or multivesicular body (MVB). IRT1 undergoes a constant cycling due to endocytosis from the plasma membrane and either degradation via the lytic pathway in the vacuole or recycling back to the plasma membrane. (A), We identified TTN5 interaction with the large cytosolic, variable region of IRT1. TTN5 and IRT1 colocalized at the plasma membrane (1). PATL2 interacts with the large cytosolic, variable region of IRT1 (IRT1vr) via its N-terminus and may be involved in α-tocopherol recruitment to the membrane, to prevent lipid peroxidation induced by IRT1-related Fe import (Hornbergs et al. 2023). TTN5 also interacts with the N-terminus of PATL2. A potential TTN5-PATL2-IRT1 complex formation is suggested (2). EHB1 interacts with the large cytosolic, variable region of IRT1 via its signature (sig) domain and inhibits IRT1 function at the plasma membrane in a Ca^2+^-dependent manner (Khan et al. 2019). TTN5 also interacted with EHB1 via the sig domain. A potential TTN5-EHB1-IRT1 complex formation is suggested (3). A possible quadruple complex of IRT1, TTN5, PATL2 and EHB1 (4). (B) We found that IRT1 colocalized with SNX1 (Ivanov et al. 2014) and TTN5 in the TGN. IRT1 and TTN5 also colocalized in MVBs. The protein representations are explained in the legend, a potential protein complex that may form, is indicated by a question mark.

### TTN5, that is part of the endomembrane system, targets IRT1vr

The protein interaction between TTN5 and IRT1 is supported by multiple findings. First, the interaction was detected in a Y2H screen, that yielded only few interaction partners, which could all be validated (Khan et al. 2019, Hornbergs et al. 2023). Second, the interaction was verified by using independent methods (targeted Y2H, BiFC, FRET-APB and Co-IP). Third, fluorescent signals of mCherry-TTN5 and IRT1-GFP reporter proteins colocalized in cells, and their localization overlapped in intracellular membrane-associated locations in cells of the root epidermis, in which both genes are expressed. Further, TTN5 was found to interact with proteins of the IRT1vr environment (Figure 5A-B). These findings altogether indicate that in plant roots, TTN5 and IRT1 indeed may interact in root epidermis cells in places related with Fe acquisition. The previously described TTN5 localization pattern in vesicles and at the plasma membrane (Mohr et al. 2024) was confirmed upon colocalization with IRT1 in this study. Hence, IRT1 localization dependent on endomembrane trafficking might be linked with the presence of TTN5 in this process.

A very interesting finding was that the mCherry-TTN5^T30N^ variant tended to colocalize more at the plasma membrane with IRT1-GFP. This could explain previous colocalization data with a MVB marker overlapping more with YFP-TTN5 and the hydrolysis impaired mutant YFP-TTN5^Q70L^ (Mohr et al. 2024) and reinforces our suggestion that TTN5 has functions in vesicle trafficking. Although a GEF is not needed for nucleotide exchange of TTN5 (Mohr et al. 2024), TTN5^T30N^ may have a conformation that facilitates binding of a GEF so that TTN5^T30N^ along with IRT1 might be retained at the plasma membrane by the unknown TTN5-interacting GEF. The presence of mCherry-TTN5 signals together with IRT1-GFP in wortmannin-induced swollen structures, probably indicating MVBs, is supporting the idea of a role in transporter degradation. MVBs are known to be the last step before vacuolar degradation (Kolb et al. 2015, Arora and Van Damme 2021). This could indicate that TTN5 presence is a critical point of IRT1 sorting fate. We therefore propose a role of TTN5 in targeting intracellular IRT1 protein (Figure 5B). In such a process, TTN5 could be present at the plasma membrane in a resting state. In the move of GTPase activation, it could be recruited to the place of action and active TTN5 would play a role in IRT1 cycling. TTN5 should predominantly bind GTP and be in an active state in the cell (Mohr et al. 2024). Further, there is support of a role of TTN5 in IRT1 recycling by the binding of TTN5 and SNX1. SNX1 is involved in the recycling of IRT1 at these sites (Ivanov et al. 2014). Hence, TTN5 can be in a functional complex with SNX1 to control the recycling of the cargo protein IRT1.

The observed roles of TTN5 fit well with the general understanding of ARF-signaling in plasma membrane protein trafficking. The auxin transporters PIN1 and PIN2 and the boron transporter BOR1, like IRT1, are polarly localized in the plasma membrane and undergo a constant degradation-recycling process (Benková et al. 2003, Takano et al. 2010, Barberon et al. 2014). Apart from vesicle formation, as described for ARF1 or SAR1, a role in this process was shown for other regulatory mechanisms of ARF-signaling through the GEFs and GAPs (Nielsen 2020). GNOM and BIG5, two ARF-GEFs, and ARF1-regulators, are described to be involved in the proper localization of the plasma membrane transporter PIN1, functioning at different regulatory steps (Geldner et al. 2003, Anders et al. 2008, Kleine-Vehn et al. 2008, Tanaka et al. 2009). In addition, another ARF GTPase ARF1A1C may not only partially colocalize with GNOM, but modified activity also resulted in impaired localization of PIN1 to the plasma membrane and also altered vacuolar targeting of PIN2 (Tanaka et al. 2014). BOR1 plasma membrane localization depends on GNOM as well (Yoshinari et al. 2021), showing a good connection between ARF-GEFs and plasma membrane protein localization.

Taken together, TTN5 that is part of the endomembrane system (Mohr et al. 2024) is linked with a transporter protein whose abundance is controlled by endomembrane trafficking. For future studies, stable coexpression of both proteins under the control of the endogenous promoter will be suitable to investigate the effect of TTN5 on IRT1 localization. Finally, due to the very low intrinsic GTP hydrolysis activity of TTN5, identification of TTN5-GAPs will shed light on TTN5 function in Fe homeostasis and potential other cellular signaling events. It will also be interesting to explore whether a membrane-localized GEF exists that can act upon TTN5. Future identification of potential TTN5-GEF and -GAP proteins will be of great interest to clarify the role of TTN5 in endomembrane trafficking with particular reference to cues controlling IRT1.

### TTN5 might play a general role in Fe nutrition

TTN5 does not only have an influence on IRT1 but also on other aspects in Fe nutrition. A partial lack of TTN5 results in decreased Fe reductase activity. The decrease in Fe reductase activity is likely caused by an effect on FRO2. It was found that IRT1 can interact with FRO2 to form a complex for a highly effective Fe uptake machinery (Martín-Barranco et al. 2020). Reduced Fe reductase activity could be due to an indirect influence of TTN5 on the complex formation or it could be directly driven by TTN5. Interestingly, both *ehb1* and *patl2* seedlings have a higher Fe reductase activity under Fe deficiency compared to wild type (Khan et al. 2019, Hornbergs et al. 2023). In case EHB1 and PATL2 also act on FRO2, this points to an opposing role of TTN5 on the Fe deficiency response in the protein complex. Gene expression analysis revealed no great impact on Fe-related genes in *ttn5-1^+/-^* roots, implying an effect of TTN5 mostly on posttranslational level.

It is a possibility that TTN5 has very different effects depending on its localization in the cell. For example, the effect on Fe reduction could be different from its effect on colocalization with IRT1 in MVBs. Different types of interaction partners were reported for the human ARL2 with connection to distinct cellular functions in tubulin folding and microtubule dynamics (Bhamidipati et al. 2000) or in mitochondria (Sharer et al. 2002). Based on these distinct locations and the associated functions of ARL2, it was proposed that ARL2 may act at a high level in signaling (Francis et al. 2016). This can be also the case in Arabidopsis. For example, TTN5 may bridge TTN5 microtubule function (Mayer et al. 1999) with IRT1 regulation.

### TTN5 interacts with EHB1, SNX1 and PATL2 and may coordinate IRT1 regulation

The close interaction of TTN5 with EHB1 was suggested by BiFC and FRET-APB and with PATL2 and SNX1 by BiFC. The interactions are further supported by deletion mutant approaches. Moreover, the interacting sites of PATL2 and EHB1 with TTN5 are similar to those for IRT1vr, supporting again the possibility that these domains and motifs are at the basis of a protein complex forming at the IRT1vr regulatory platform.

Interaction with the IRT1 regulators EHB1, PATL2 and SNX1 suggests that TTN5 could not only be linked with Fe nutrition but also the associated Ca^2+^- and ROS-signaling events (Khan et al. 2019, Hornbergs et al. 2023). EHB1 is additionally interesting as a member of the CAR family. Several CAR proteins can initiate Ca^2+^-dependent membrane curvature, which may provide a direct link to vesicle trafficking. Interestingly, Ca^2+^ is bound by EHB1 *via* its C2 domain, which is also a common feature of class III ARF-GAPs (Vernoud et al. 2003, Knauer et al. 2011). Indeed, EHB1 is directly linked to ARF signaling in an antagonistic interaction with ARF-GAP AGD12 in gravitropic bending (Dümmer et al. 2016, Rath et al. 2020). Further research needs to investigate whether TTN5 can interact with the C2-domain containing ARF-GAPs AGD11-AGD13 as well. Additionally, crystallization experiments revealed a potential Ca^2+^-binding site between HsARF6 and its GAP ASAP3. The GTP hydrolysis was stimulated in the presence of Ca^2+^ proposing a linkage between Ca^2+^- and ARF-signaling (Ismail et al. 2010). We therefore consider the possibility of an interaction between TTN5 and EHB1 in a Ca^2+^-dependent manner. PATL2 may interact with IRT1 to prevent lipid peroxidation stress (Hornbergs et al. 2023). Prolonged Fe deficiency leads to an increase in hydrogen peroxide concentration as well as an activity increase of catalase CAT2 (Le et al. 2015, von der Mark et al. 2020, Gratz et al. 2021). Some catalases are present in the PATL2 interactome in roots (Hornbergs et al. 2023). Future studies may address whether TTN5 is connected with oxidative stress and Ca^2+^ signaling.

SNX1 is a positive regulator for IRT1 recycling to the plasma membrane in the sorting endosomes (Ivanov et al. 2014) and for endosomal recycling of the auxin efflux carrier PIN2 (Jaillais et al. 2006). SNX1 is therefore part of the recycling of specific plasma membrane proteins. Interestingly, deficiency of BLOS1 leads to accumulation of PIN1 and PIN2 at the plasma membrane (Cui et al. 2010). BLOS1 seems to be an ortholog of the mammalian biogenesis of lysosome-related organelles complex 1 (BLOC-1), which is responsible for vesicle transport from endosomes to lysosomes (Li et al. 2007, Raposo et al. 2007). SNX1 can interact with both BLOS1 and BLOS2 and could therefore also play a role in vesicle-mediated transport through these interactions (Cui et al. 2010, Heucken and Ivanov 2018). SNX1 possesses a BAR domain which is also typical for class I ARF-GAPs (Vernoud et al. 2003, Heucken and Ivanov 2018). A connection between the SNX1 BAR domain and an ARF6-GEF was identified in mouse by their interaction. This interaction was associated with a positive effect of ARF6 function (Fukaya et al. 2014). Colocalization analysis revealed overlapping expression in endosomes, which fits known SNX1 localization in Arabidopsis (Fukaya et al. 2014, Ivanov et al. 2014), this could be a possible interconnection between ARF-signaling and SNX1. Here, we identified an interaction between TTN5 and SNX1, which could be the critical link in the decision between SNX1-dependent recycling and degradation of IRT1. SNX1 could be needed for a potential TTN5-GEF and subsequent TTN5 recruiting to membranes. Based on the opposing roles of EHB1 and SNX1 in IRT1 regulation and their connection to TTN5, we suggest a coordinating role of TTN5 in transporting IRT1 between plasma membrane and endosomes by interacting with peripheral membrane proteins such as EHB1, PATL2 and SNX1. The findings that some ARF-GAPs have a C2 or BAR domain, and that SNX1 interacts with an ARF-GEF strengthen the interest in the identification of TTN5-GEFs and -GAPs.

Further research has to focus on physiological aspects in either combined knockout mutants or tagged-protein lines to arrange and decipher the IRT1 regulatory network. An intriguing open question is whether a large IRT1vr-EHB1-PATL2-TTN5-SNX1 complex is assembled. If so, the next question is how, when and in which order the individual or all components of the IRT1vr complex are assembled. Another question is what the structural requirements for proper protein conformations and membrane interfaces are (Figure 5A, B).

### Conclusion and perspectives

This study provides hints that the ARF-like GTPase TTN5 contributes to endocytosis and vesicle trafficking of IRT1 by way of a protein complex at IRTvr comprising TTN5 and other IRT1 interactors. Evidence that small GTPases can bind with plasma membrane and peripheral membrane proteins is instrumental to understand how membrane proteins are controlled by vesicular trafficking components in plants. An intriguing question is whether TTN5 also targets other transmembrane cargo proteins. Future work may uncover additional functions of TTN5 or of other GTPase protein complexes of this kind and determine the requirements and order of events for complex assembly leading to vesicular trafficking.

## Material & Methods

### Yeast two-hybrid (Y2H)

The yeast two-hybrid screen for identification of IRT1vr interactors was performed previously and is described in Khan et al. (2019). Targeted Y2H interactions were tested using the GAL4 transcription factor system with histidine as selection marker. *TTN5* coding sequence was amplified using TITAN5 nter B1 and TITAN5 stop B2 primers (for primer sequences, see Supplemental Table S1). Obtained fragment was cloned into pDONR207 *via* Gateway BP reaction (Life Technologies) with following LR reactions into pGBKT7-GW (binding domain, BD) or pACT2-GW (activation domain, AD). Both vectors were a kind gift from Dr. Yves Jacob. The yeast strain AH109 was cotransformed with the corresponding AD- and BD-tagged expression vectors. The combination pACT2:SNX1 and pGBKT7:SNX1 (Pourcher et al. 2010) was used as a positive control. Negative controls were cotransformations with the respective non-recombined pACT2-GW or pGBKT7-GW. Drop test was performed with SD -Leu, -Trp-cultures at an OD_600_ of 0.1 and three additional 1:10 dilution steps (10^1^-10^4^) on SD -Leu, -Trp- and SD -Leu, -Trp, -His +0,5 mM 3-amino-1,2,4-triazole-plates. One of each plate was incubated either at 30°C or at room temperature, wrapped in aluminum foil for up to two weeks.

### Nicotiana benthamiana leaf infiltration

*Nicotiana benthamiana* plants were grown on soil for 2-4 weeks in a greenhouse facility under long day conditions (16 hours light, 8 hours darkness, 20°C). *N. benthamiana* leaf infiltration was performed with the Agrobacterium (*Rhizobium radiobacter*) strain C58 (GV3101) carrying the respective constructs for confocal microscopy. Agrobacteria cultures were grown overnight at 28°C, centrifuged for 5 min at 4°C at 5000*g*, resuspended in infiltration solution (5% sucrose, a pinch of glucose, 0.01% Silwet Gold, 150 µM acetosyringone) and incubated for 1 hour at room temperature. Bacterial suspension was set to an OD_600_=0.4 and infiltrated into the abaxial side of *N. benthamiana* leaves. Expression of pABind and pMDC7 (Curtis and Grossniklaus 2003, Bleckmann et al. 2010) constructs was induced with a β-estradiol solution (20 µM β-estradiol, 0.1% Tween 20) 16 hours before imaging.

### Bimolecular Fluorescence Complementation (BiFC)

BiFC was used to study protein interaction of TTN5 and the respective GTPase TTN5^T30N^ and TTN5^Q70L^ variants with other proteins in plant cells. At first, entry clones were generated. *TTN5*, *TTN5^T30N^* and *TTN5^Q70L^* coding sequences with stop codon were amplified using the primers TITAN5 n-ter B1 and TITAN5 stop B4 (Supplemental Table S1). *IRT1vr* coding sequence with stop codon was amplified using the primers I1L1B3 and I1LB2. Amplifying full-length *EHB1* the primer pair EHB1 n-ter B3 and EHB1 stop B2 was used. Amplification of *EHB1Δsig* is described in Khan et al. 2019. *SNX1* coding sequence without stop codon for C-terminal fusion was amplified using the primer SNX1 B3 and SNX1 ns B2. Cloning of SNX1 deletion mutants SNX1PX and SNX1BAR was done using the primers SNX1 B3 and PXSNX1 rev and BARSNX1 fwd and SNX1 ns B2 respectively. The used PATL2 constructs are described in Hornbergs et al. (2023). The amplified PCR products were cloned *via* Gateway BP reaction (Life Technologies) into pDONR221-P1P4 (Invitrogen) or pDONR221-P3P2 (Invitrogen).

The obtained constructs were then used to prepare final recombinant destination vectors in an LR reaction (Life Technologies) for cloning into pBiFCt-2in1 vectors (Grefen and Blatt 2012). Agrobacteria were transformed with correct constructs and used for *N. benthamiana* leaf infiltration. After 48 hours, the mRFP expression control signal and YFP signals were detected by fluorescent microscopy (LSM 780, Zeiss) with a 40x C-Apochromat water immersion objective. YFP constructs were detected at 491-560 nm after exciting at 488 nm and mCherry fluorescence was excited at 561 nm and emission detected at 570-633 nm.

The BiFC constructs were tested in three independent replicates with three infiltrated leaves each. The vectors pBiFC-2in1-NN and pBiFC-2in1-CN were kindly provided by Dr. Christopher Grefen, Tübingen, Germany.

### Förster-Resonance-Energy Transfer Acceptor Photo Bleaching (FRET-APB)

FRET-APB was used to verify protein-protein interactions of TTN5 TTN5^T30N^, and TTN5^Q70L^. The coding sequences without stop codon were amplified with the primers TITAN5 B1 and TITAN5 ns B2, and cloned into the pDONR207 (Invitrogen, BP reaction, Life Technologies). Cloning of pDONR207:IRT1vr and pDONR207:EHB1 is described in Khan et al. (2019). The obtained constructs were used for LR reactions (Life Technologies) with the pABind-GFP, pABind-mCherry and pABind-FRET for C-terminal tagging (Bleckmann et al. 2010). N-terminal GFP-TTN5 constructs were cloned by overlap extension PCR simultaneously with mCherry-TTN5 constructs (check paragraph ‘Subcellular localization of fluorescent protein fusions’). In the first step the primer pairs GFP B1 with GFP R ns BIND and GFP to TTN5 with TITAN5 stop B2 were used with pABind-FRET (Bleckmann et al. 2010) as template for the fluorescent protein coding sequence. The final construct was obtained with the primer pair GFP B1 with TITAN5 stop B2. Constructs were cloned into pDONR207 for further cloning into pMDC7 *via* LR reactions. Agrobacteria were transformed and used for *N. benthamiana* leaf infiltration. FRET-APB was performed using laser-scanning confocal microscopy (LSM 780, Zeiss) with a 40x C-Apochromat water immersion objective. GFP was excited at 488 nm and detected at 491-560 nm and mCherry fluorescence was excited at 561 nm and detected at 570-633 nm. GFP and mCherry channels were recorded for five scans. The mCherry signal was then bleached with 70 iterations of maximum laser intensity in a specific region of interest (ROI) which was set to the nucleus. Both channels were detected for additional 20 post-bleaching scans. FRET-APB measurements were performed with a minimum of 10 repetitions (n ≥ 10).

### Arabidopsis plant material

The Arabidopsis *ttn5-1* mutant was previously described (McElver et al. 2000). Heterozygous seedlings were selected by PCR on gDNA using the primer TTN5 intron1 fwd and pDAP101 LB1 (Supplemental Table S1). pro35S::YFP-TTN5 and pro35S::HA_3_-TTN5 lines were previously described in (Mohr et al. 2024) and proIRT1::IRT1-mCitrine/*irt1-1* in (Dubeaux et al. 2018). *irt1-1* (Ws) was characterized in (Vert et al. 2002) and *irt1-1* (Col-0, SALK_054554) in Fukao et al. (2011).

Reporter lines were constructed as follows: proIRT1::IRT1-mCherry, with the mCherry at position 80 in the first cytoplasmic loop, was cloned by overlap-extension PCR. *mCherry* was amplified from pABind:mCherry (Bleckmann et al. 2010) using primer pair link mCh F and mCh ns link R. IRT1 halves were amplified using IRT1 B1 and IRT1Q80 link R and link IRT1Q80 F and IRT1 stop B2. In a last PCR IRT1-mCherry was amplified with primer pair IRT1 B1 and IRT1 stop B2. BP and LR reaction (Life Technologies) were performed as described above with pIM5 as destination vector. The Gateway-suitable plasmid pIM5 was created by exchanging pro35S from pMCD32 with proIRT1 using AQUA cloning (Beyer et al. 2015). The IRT1 promoter was amplified from gDNA using primer pair pIM5-pIRT1 F and pIM5-pIRT1 R with pMDC32-complementary overhangs and the restriction sites HindIII and KpnI respectively. IRT1-GFP construct, with the GFP at position 173 in the variable, cytoplasmic loop, was cloned by overlap-extension PCR. *GFP* was amplified from pABind:GFP (Bleckmann et al. 2010) using primer pair IG173 F and GI173 R. IRT1 halves were amplified using IRT1 B1 and IG173 R and GI173 R and IRT1 stop B2. In a last PCR IRT1-GFP was amplified with primer pair IRT1 B1 and IRT1 stop B2. BP and LR reaction were performed as described above with pMDC32 (Curtis and Grossniklaus 2003) as destination vector. Agrobacterium cultures containing the plant protein expression vectors, generated as described above, were used for floral dip transformation of *A. thaliana*. Plants were multiplied, PCR-genotyped using specific primer pairs (Supplemental Table S1) and selected according to standard procedures.

### Co-Immunoprecipitation experiments/pull-down

Immunoprecipitation (IP) was performed with Arabidopsis seedlings grown for 10 days under Fe-deficient conditions. The proIRT1::IRT1-mCherry in either HA_3_-TTN5 or WT background were grown and analyzed in parallel. Approximately 60 seedlings were flash-frozen in liquid nitrogen. and grinded using the Retsch MM200 with cooled grinding jars in the presence of two metal balls for 2 min. Grinded material was solubilized in 1 ml IP Buffer (50 mM Tris-HCl, 150 mM NaCl, 1 mM EDTA, 1% Triton X-100 and cOmpete Mini Protease Inhibitor, pH 8). Solution was incubated for 10 min at room temperature followed by a 10 min centrifugation at 4°C and 17.000*g* (max. speed). The supernatant was transferred into a fresh 1.5 ml reaction tube. The centrifugation step was repeated until all cellular debris were removed. 50 µl were taken as Crude Extract (CE) aliquot. 25 µl anti-HA magnetic beads (Thermo Scientific, Pierce) were added to the solution and reaction was incubated overnight at 4°C while rotating. A magnet was used to collect the beads at the side of the reaction tube and solution was transferred to a fresh 1.5 ml reaction tube (Flow through). 1 ml IP buffer was added to wash beads (Wash I). Collection of the supernatant and washing procedure was repeated for a total number of two washing steps. IP was performed in two replicates each (n = 2)

### Arabidopsis physiological Fe deficiency growth experiments

Arabidopsis seeds were sterilized with sodium hypochlorite solution (6% Sodium hypochlorite and 0.1% Triton X-100) and stored 24 hours at 4°C for stratification. Seedlings were grown upright on Hoagland media [1.5 mM Ca(NO_3_)_2_, 0.5 mM KH_2_PO_4_, 1.25 mM KNO_3_, 0.75 mM MgSO_4_, 1.5 µM CuSO_4_, 50 µM H_3_BO_3_, 50 µM KCl, 10 µM MnSO_4_, 0.075 µM (NH_4_)_6_Mo_7_O_24_, 2 µM ZnSO_4_ and 1% sucrose, pH 5.8, supplemented with 1.4% Plant agar (Duchefa)] with sufficient (50 μM FeNaEDTA, +Fe) or deficient (0 μM FeNaEDTA, -Fe) Fe supply in growth chambers (CLF Plant Climatics) under long day condition (16 hours light at 21°C, 8 hours darkness at 19°C). Seedlings were grown on Fe-sufficient or -deficient media for 6 or 10 days or in a two-week growth system with plants growing 14 days on Fe-sufficient media and then transferred to either fresh Fe-sufficient or Fe-deficient media for additional three days.

### Root length

For root length comparison plants were growing for 10 days in Fe-sufficient or -deficient conditions photographed including a scale in the picture. Root length were measured from root tip to hypocotyl using the JMicro Vision program (Roduit, N. JMicroVision: Image analysis toolbox for measuring and quantifying components of high-definition images. Version 1.3.4. https://jmicrovision.github.io) (+Fe: WT n = 19; *ttn5-1* n = 28; -Fe: WT n = 16; *ttn5-1* n = 32).

### Chlorophyll content

For chlorophyll content comparison plants were grown in the two-week system in Fe-sufficient or -deficient conditions. Chlorophyll content was measured by SPAD values using SPAD-502plus (Konica Minolta Optics, 2012) with up to ten biological replicates (+Fe: n = 10; -Fe: n = 9). Chlorophyll content was determined in three individual experiments.

### *ttn5-1* phenotyping on alkaline calcareous soil

Sterilized seeds were placed on Fe-sufficient Hoagland plates in three rows with 12-15 seeds, stratified at 4°C in darkness for three days and then (day of sowing [DAS]) grown upright for nine days. Sixteen wildtype and sixteen *ttn5-1^+/-^*seedlings were selected for transfer to on two soil conditions, control and alkaline calcareous soil (ACS). Control condition consisted of peat-based soil (Floraton 1, Floragard, Oldenburg, Germany), 20 g vermiculite (Agrivermiculite Floragard, Oldenburg, Germany) and 400 ml distilled water per liter dry soil. The ACs soil additionally contained 8 g CaCO_3_ (AppliChem, Darmstadt, Germany) and 4 g NaHCO_3_ (Fisher Scientific, Hampton, USA). The pH in water (1:10 soil:water) was 6.2 for the control soil and 8.0 for the ACS soil 30 minutes after mixture. The soil was filled into 7 cm*7 cm*6.5 cm pots (Pöppelmann GmbH & Co. KG Kunststoffwerk–Werkzeugbau, Lohne Germany) and afterwards 7.5 cm *7.5 cm large squares of ultramarine blue plotter foil (Europe Warehouse GmbH & Co. KG, Wuppertal, Germany) were stuck on the pots. Seedling were then planted into round holes in the foil with 1 cm diameter. Eight plants were grown per soil and condition. After one week, the 1 cm large holes were closed around the seedlings.

Plants were watered for one week with deionised water. Then control plants were watered with deionised water and ACS plants with 20 g/l NaHCO_3_ (second week) and 25 g/l NaHCO_3_ (following weeks). 28 days after sowing the plants were scanned with a MicroScan with PlantEye (Phenospex, Heerlen, The Netherlands). It is a multispectral scanner determining positions and color of points using a laser and four additional wavelengths (red: 624-634 nm, green: 530-540 nm, blue: 465-485 nm, near-infrared: 720-750 nm). By filtering out the blue color (hue 200-360) form the images the background around the plants was removed from the measurements. Each plant was scanned twice and the average of the technical replicates was calculated. The parameter “3D Leaf Area” and “Normalized Differential Vegetation Index” (NDVI) average were determined with the PlantEye MicroScan (Phenospex, Belgium) according to the instruction manual, corresponding to an approximation of aerial plant biomass and chlorophyll content. Photos of the plants were acquired with a camera (α 600, Sony, Tokyo, Japan).

### Gene expression analysis by RT-qPCR

For gene expression analysis, plants were grown in the two-week system. Total RNA was isolated from roots using the RNeasy Mini Kit (Qiagen) and cDNA synthesis was performed with the RevertAid RT Reverse Transcription Kit (ThermoFisher Scientific). The CFX96 Touch^TM^Real-Time PCR Detection System (Bio-Rad) was used for RT-qPCR performance. Data processing was done with Bio-Rad CFX Manager^TM^ (version3.1) software. Mass standard curve analysis was used for determination of the absolute gene expression and the elongation factor *EF1Bα* expression served as a reference for normalization. RT-qPCR was performed in three biological replicates (n = 3) and two technical replicates each. All primer pairs used in this study are listed in Supplemental Table S1.

### Fe reductase activity assay

Fe reductase activity assay was performed as described in (Gratz et al. 2019). For Fe reductase activity plants were grown in the two-week system in Fe-sufficient or -deficient conditions. Two plants per replicate were washed in 100 mM Ca(NO_3_)_2_, then incubated for 1 hour in the dark in 1.5 ml Fe reductase solution (300 mM ferrozine and 100 mM FeNaEDTA). The Fe reductase activity-dependent color change was measured at 562 nm using Infinite 200® PRO (Tecan) plate reader. Activity calculation was done using the ferrozine extinction coefficient ε = 28.6 mM^-1^*cm^-1^ and was normalized to root weight. The assay was performed with a minimum of three replicates (n ≥ 3), each consisting of a pool of two plants. Fe reductase activity was determined in three individual experiments.

### Subcellular localization of fluorescent protein fusions and microscopy

The creation of TTN5^T30N^ and TTN5^Q70L^ constructs by introduced point mutations and generation of pro35S::YFP-TTN5 is described in Mohr et al. (2024). mCherry-tagged constructs were created by overlap-extension PCR. Fluorescent protein was amplified from pABind:mCherry (Bleckmann et al. 2010) using primer pair mCherry B1 and mCh R ns BIND. *TTN5*, *TTN5^T30N^* and *TTN5^Q70L^* CDS were amplified with a 20 bp overlap of mCherry using mCh to TTN5 and TITAN5 stop B2. In a third PCR mCherry-tagged TTN5 constructs were generated with the mCherry B1 and TITAN5 stop B2 primer pair. BP and LR reaction were performed as described above. As destination vector the XVE-driven β-estradiol inducible vector pMDC7 (Curtis and Grossniklaus 2003) was chosen. Cloning of IRT1-GFP is described above in the section ‘Arabidopsis plant material’. IRT1-mCherry construct is described in Ivanov et al. (2014) and proIRT1::IRT1-mCitrine/*irt1-1* in Dubeaux et al. (2018). Agrobacteria were transformed with the obtained constructs and tested by *N. benthamiana* leaf infiltration after two days of expression.

Localization studies were carried out by laser-scanning confocal microscopy (LSM 780, Zeiss) with a 40x C-Apochromat water immersion objective. YFP constructs were excited at 488 nm and detected at 491-560 nm. mCherry or FM4-64 fluorescence was excited at 561 nm and detected at 570-633 nm.

Wortmannin (10 µM, Sigma-Aldrich) and plasma membrane dye FM4-64 (165 µM, ThermoFisher Scientific) were infiltrated into *N. benthamiana* leaves. FM4-64 was detected after 5 min incubation and Wortmannin was incubated for 25 min before checking the treatment effect.

### JACoP based colocalization analysis

Colocalization analysis was performed with the ImageJ (Schneider et al. 2012) Plugin Just Another Colocalization Plugin (JACoP) (Bolte and Cordelières 2006). A comparison of Pearson’s and Overlap coefficients was done. Object-based analysis was performed for spotted-structures, adapted by Ivanov et al. (2014). Percentage of colocalization for both channels was calculated based on distance between geometrical centers of signals. Analysis was conducted in three replicates each (n = 3).

### Statistical analysis

One-way ANOVA was used for statistical analysis and performed in OriginPro 2019. Fisher LSD or Tukey was chosen as post-hoc test with p < 0.05.

### ACCESSION NUMBERS

Sequence data from this article can be found in the TAIR and GenBank data libraries under accession numbers: *FRO2* (TAIR: AT1G01580), *EHB1* (TAIR: AT1G70800), *IRT1* (TAIR: AT4G19690), *PATL2* (TAIR: AT1G22530), *TTN5* (TAIR: AT2G18390), and *SNX1* (TAIR: AT5G06140). Raw microscopic image data are uploaded and will be available upon publication of this manuscript via BioImage Archive (https://www.ebi.ac.uk/bioimage-archive/).

## Acknowledgements

We thank Gintaute Matthäi and Elke Wieneke for excellent technical assistance. We thank Ksenia Trofimov for microscopic help and advice, and Rubek Merina Basgaran for help with protein interaction studies. We are thankful for the excellent assistance from Stefanie Weidtkamp-Peters and Sebastian Hänsch, members of the Center for Advanced Imaging (CAi) at the Heinrich Heine University. This work was supported by the Deutsche Forschungsgemeinschaft (DFG, German Research Foundation), SFB 1208 project B05 to P.B.. This work received funding by Deutsche Forschungsgemeinschaft (DFG, German Research Foundation) under Germanýs Excellence Strategy – EXC-2048/1 – project ID 390686111 and – TRR 341/1 - Project-ID 456082119. Funding for instrumentation: Zeiss LSM780: DFG-INST 208/551-1 FUGG.

## Abbreviations

AD: activation domain

Arabidopsis: *Arabidopsis thaliana*

ARF-like: ARL ADP-ribosylation factor-like

BD: binding domain

EE: early endosomes

EHB1: ENHANCED BENDING 1

FRET-APB: Förster Resonance Energy Transfer-Acceptor Photobleaching

GAP: GTPase-activating protein

GEF: guanine nucleotide exchange factor

MVB: multivesicular body

PATL2: PATELLIN 2

ROI: region of interest

SNX1: SORTING NEXIN 1

TGN: *trans*-Golgi network

TTN5: TITAN 5

Y2H: yeast two-hybrid

ZIP: ZRT/IRT-LIKE PROTEIN

---

**Supplemental Figure S1.**
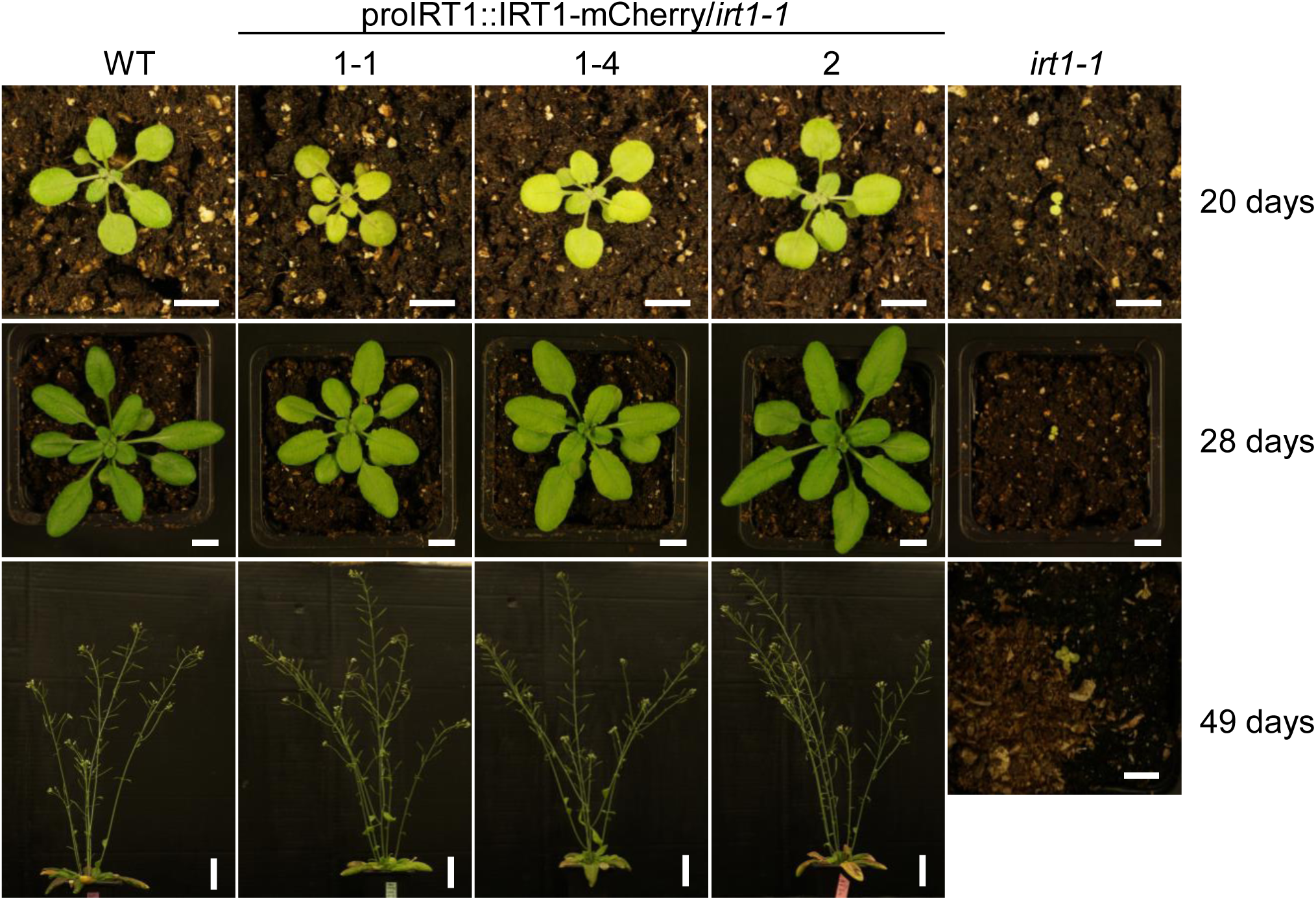
IRT1Q80-mCherry complements *irt1-1* knockout phenotypes. Complementation of the rosette growth defect, leaf chlorosis and reproductive capacity of *irt1-1* (SALK_054554) by IRT1-mCherry. Seedlings of wild type (WT), proIRT1::IRT1-mCherry/*irt1-1* (briefly IRT1-mCherry) and *irt1-1* were grown on soil. Rosettes were imaged after 20, 28 and 49 days. IRT1-mCherry lines complemented both the defect in rosette growth, leaf chlorosis, and silique formation of the *irt1-1* mutant and were comparable to the wild type. Scale bar horizontal: 1 cm, vertical: 5 cm.

**Supplemental Figure S2.**
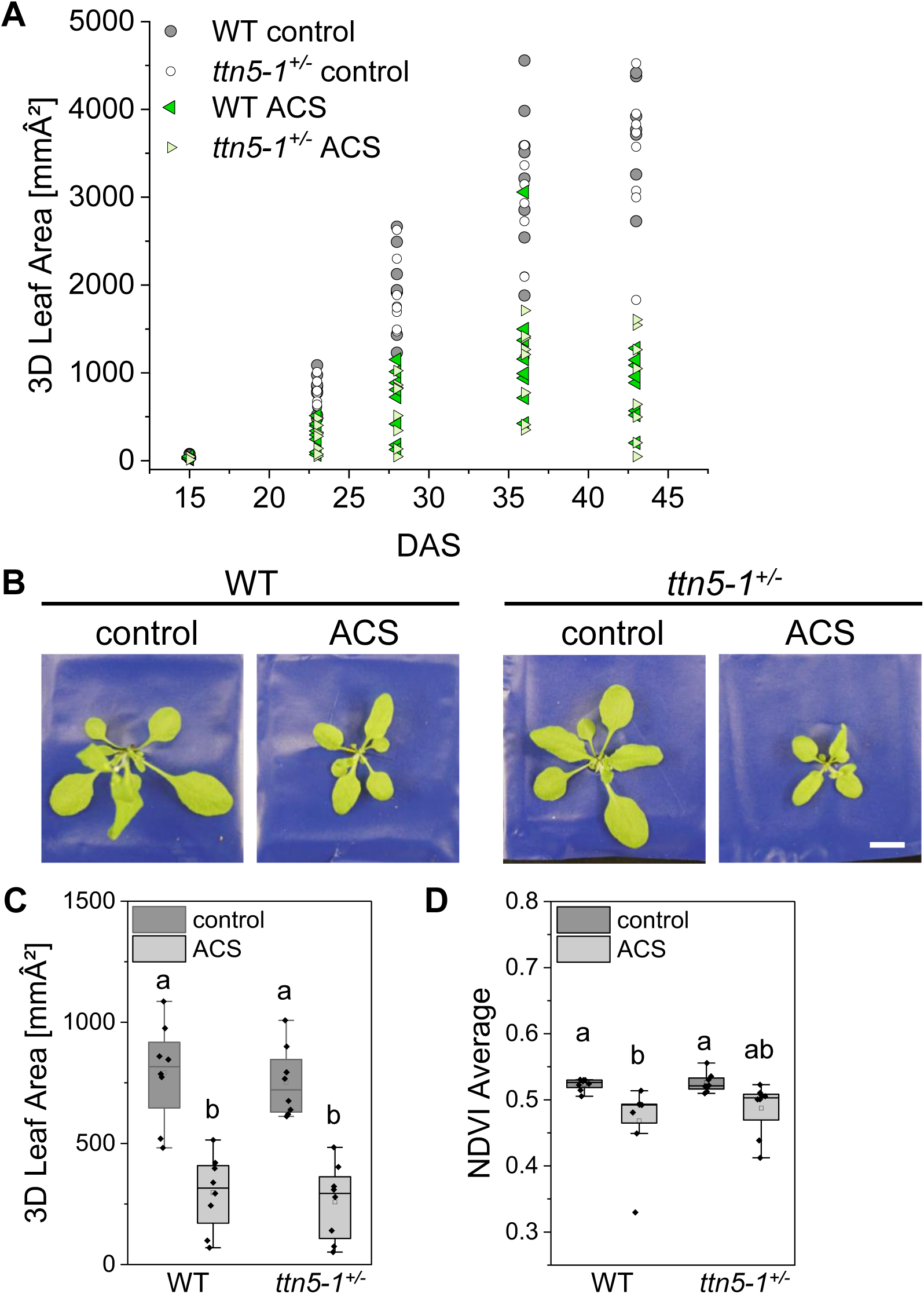
Alkaline calcareous soil similarly impacts growth of *ttn5-1^+/-^* and wild type plants. Heterozygous *ttn5-1^+/-^* and their wild type sibling seedlings were grown in comparison on +Fe Hoagland plates for 9 days and then transferred to alkaline, calcareous soil (ACS) or control conditions. Phenotypical analysis was conducted 15, 23, 28, 35 and 43 days after sowing using PlantEye (Phenospex). (A), Growth of wild type (WT; dark colors) and *ttn5-1^+/-^* (bright colors) plants on both, control (circle) and ACS (triangle) soil conditions determined by 3D leaf area. The parameter “3D Leaf Area” represents the area of the surface of the whole scanned object visible from straight above, including rosettes, stems and inflorescences. ACS condition led to a reduced 3D leaf area for both wild type and *ttn5-1^+/-^*. (B-D), Detailed phenotypical analysis of WT and *ttn5-1^+/-^* after 23 days. (B) Images of WT and *ttn5-1^+/-^* on both soil conditions. ACS condition led to a reduced rosette size for both. Scale bar 1 cm. (C), PlantEye analysis confirmed a significant reduced 3D leaf area on ACS compared to control conditions for both wild type (WT) and *ttn5-1^+/-^* plants in a similar amount. The box represents the 25–75^th^ percentiles, and the median is indicated. The whiskers show the 5^th^ and 95^th^ percentiles. (D), Normalized difference vegetation index (NDVI) was used as a correlation factor for photosynthetic efficiency. For the NDVI, the reflectance of red (RED, 624-634 nm) and near-infrared (NIR, 720-750 nm) light is determined at all points of the scanned object, that are reached by the light. The NDVI is then calculated as (NIR – RED) / (NIR + RED) for each point and the average over the object is determined. WT plants grown in ACS condition had a significant reduced NDVI compared to control soil while this effect was less the case for *ttn5-1^+/-^* plants. The experiment was performed with eight biological replicates (n = 8). The box represents the 25–75^th^ percentiles, and the median is indicated. The whiskers show the 5^th^ and 95^th^ percentiles. One-way ANOVA with Tukey post-hoc test was performed. Different letters indicate statistical significance (p < 0.05).

**Supplemental Figure S3.**
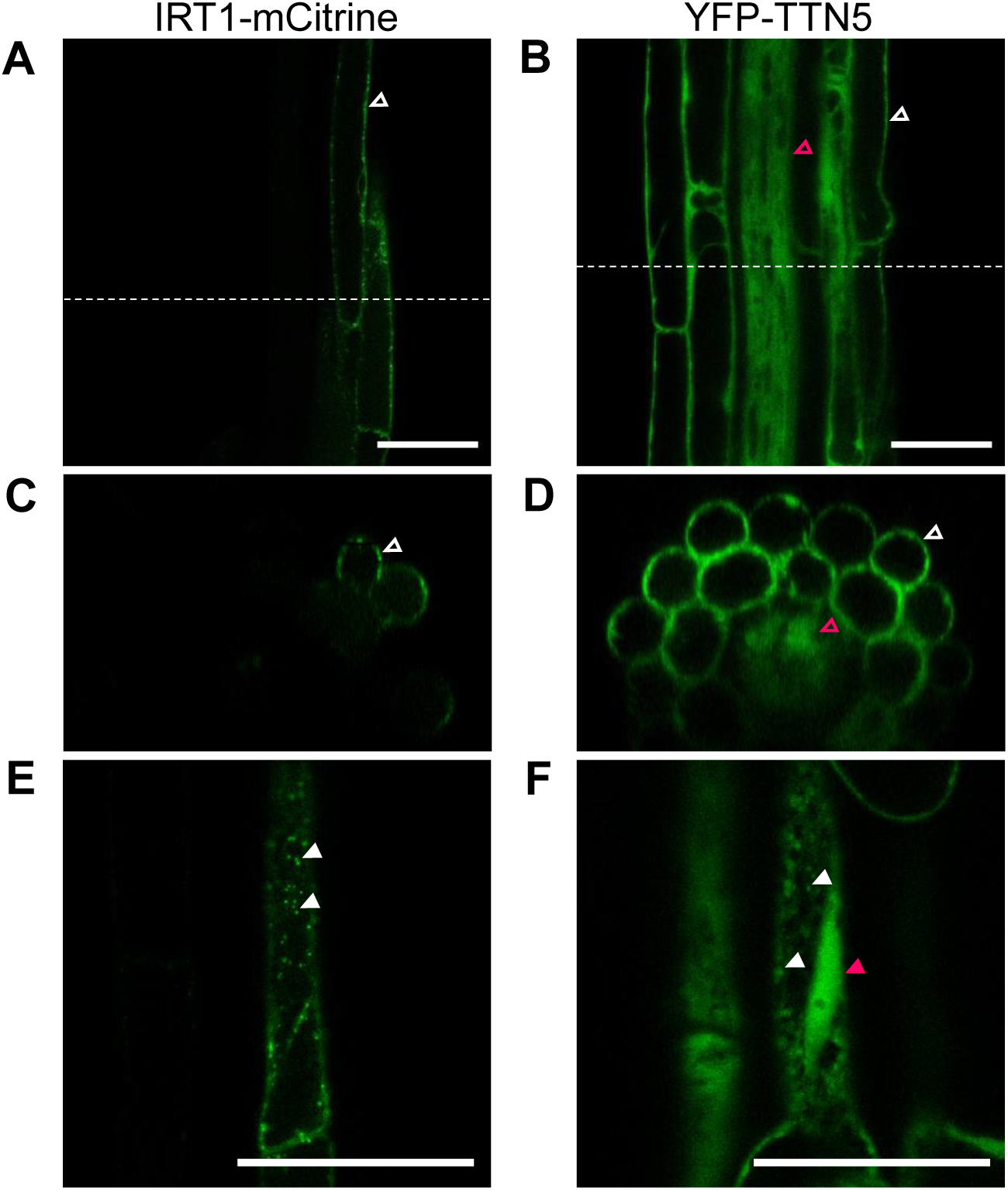
Localization of IRT1 and TTN5 fluorescence fusion proteins in the Arabidopsis root elongation zone. (A, C, E) IRT1-mCitrine localization in proIRT1::IRT1-mCitrine/*irt1-1* seedling roots (line previously described in Dubeaux et al. (2018), Hornbergs et al. (2023)) in comparison with (B, D, F) YFP-TTN5 localization in pro35S::YFP-TTN5/WT seedling roots (line previously described in Mohr et al. (2024)) in the root elongation zone. Plants were grown on plates side by side. (A, B), Longitudinal confocal sections through the stele. (A) IRT1-mCitrine was present at the plasma membrane of epidermal cells (empty white arrowhead). (B) Fluorescent signal of YFP-TTN5 roots was localized at the plasma membrane of epidermal cells and other cell types, as well as in the stele (empty magenta arrowhead). Dotted white lines indicate the position of the cross sections shown in C and D. (C-D), Cross confocal sections highlighting (C) IRT1-mCitrine in epidermal cells compared to (D) nearly ubiquitously localized fluorescent signal in YFP-TTN5 roots. (E-F), Longitudinal section through the epidermis regions. Next to plasma membrane, (E) IRT1-mCitrine and (F) YFP-TTN5 were present in intracellular spots (filled white arrowheads). YFP-TTN5 was also detected in the nucleus (filled magenta arrowhead). The experiment was repeated in a minimum of three times (n ≥ 3). Scale bar 10 µm.

**Supplemental Figure S4.**
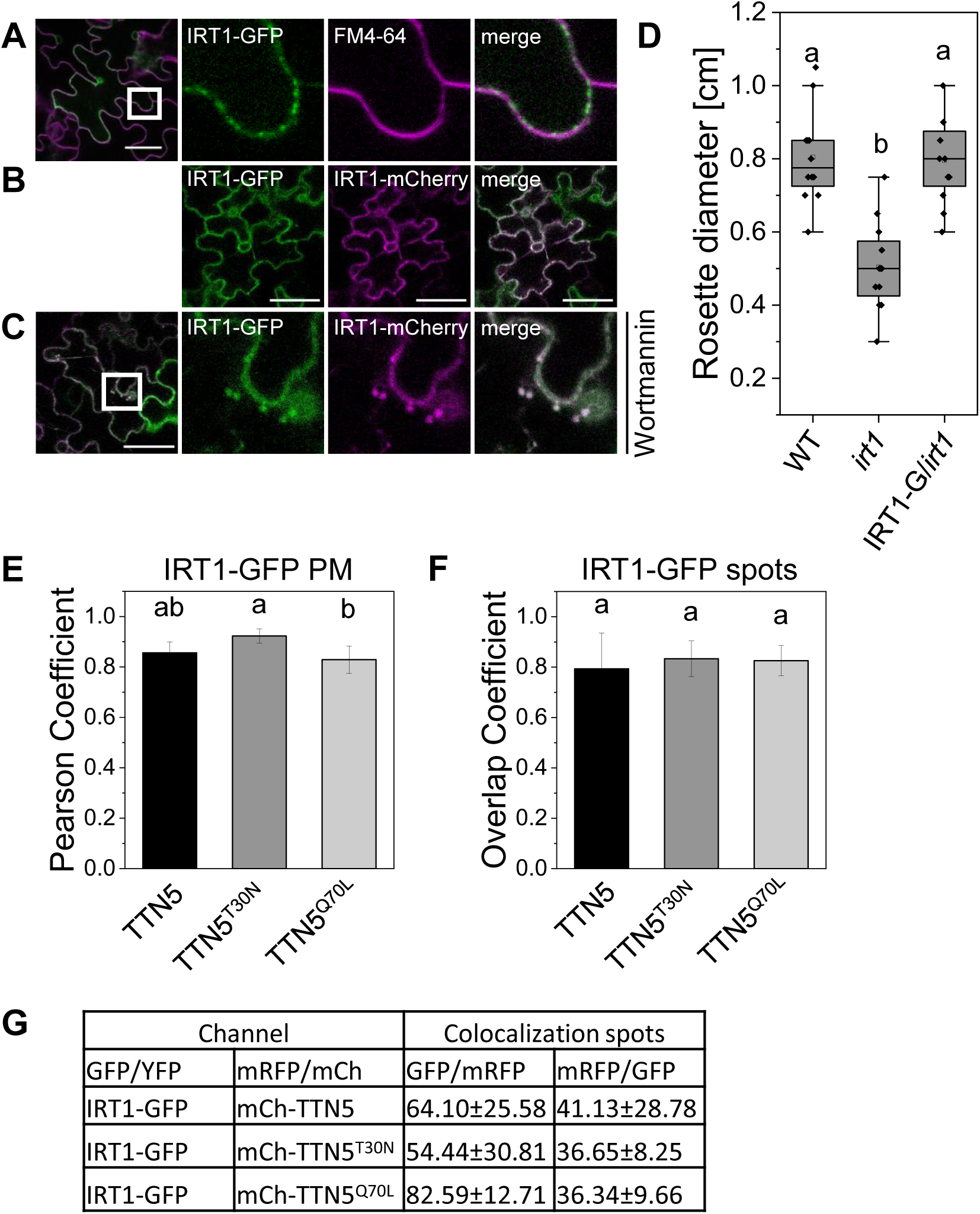
IRT1 colocalization analysis at the plasma membrane and in intracellular spots. (A), Plasma membrane dye FM4-64 indicated IRT1-GFP localization at the plasma membrane. IRT1-GFP is present in a spotted structure, indicating regions of higher transporter accumulation. (B-C), IRT1-GFP colocalized with IRT1-mCherry (Ivanov et al. 2014) (B) at the plasma membrane (PM) and (C) in swollen intracellular spots after wortmannin treatment (10 µM), confirming correct localization of IRT1-GFP. The experiments were repeated in a minimum of three times (n ≥ 3). Scale bar 50 µm. (D), Complementation of rosette growth defect of *irt1-1* by IRT1-GFP. Rosette sizes of *irt1-1* and pro35S::IRT1-GFP/*irt1-1* were measured 14 days after sowing. IRT1-GFP complemented the growth defect of *irt1-1*. The experiment was conducted once with twelve biological replicates (n = 12). The box represents the 25–75^th^ percentiles, and the median is indicated. The whiskers show the 5^th^ and 95^th^ percentiles. (E-G), JACoP-based colocalization analysis of mCherry-TTN5 and variants with IRT1-GFP (Bolte and Cordelières, 2006) performed with ImageJ (Schneider et al. 2012). See also Figure 3A-H. (E-F), Comparison of (from left to right) Overlap coefficients for IRT1-GFP with mCherry-TTN5, mCherry-TTN5^T30N^ and mCherry-TTN5^Q70L^ (E) at the plasma membrane and (F) in vesicle-like structures. (G), Object-based analysis was performed for spotted-structures based on distance between geometrical centers of signals. Analyses were conducted in three replicates each (n = 3). One-way ANOVA with Fisher-LSD post-hoc test was performed. Different letters indicate statistical significance (p < 0.05).

**Supplemental Figure S5.**
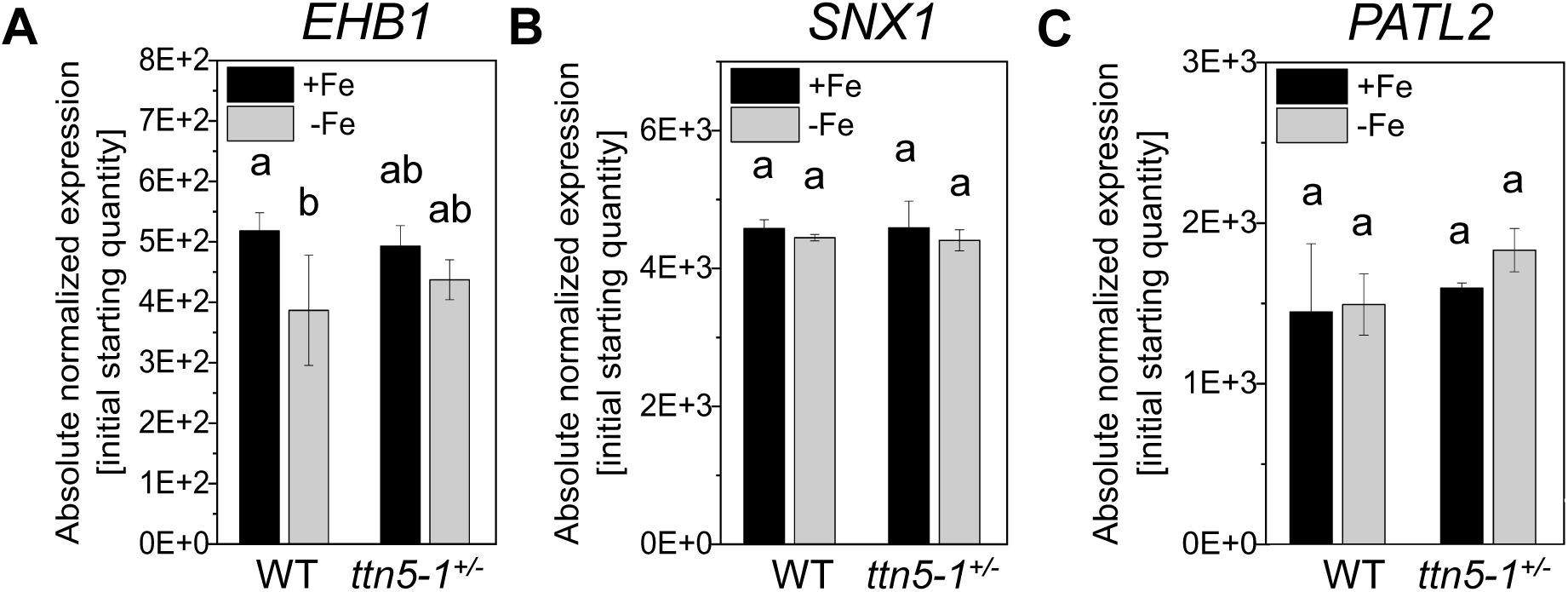
Gene expression of *EHB1*, *SNX1* and *PATL2* is not affected in *ttn5-1^+/-^* plants. Gene expression data by RT-qPCR of *EHB1*, *SNX1* and *PATL2* in *ttn5-1^+/-^* roots compared to wild type sibling roots. Plants were grown in the two-week system in Fe-sufficient (+Fe, black bars) or -deficient (-Fe, grey bars) conditions. See also Figure 2D-F. (A), *EHB1* expression was down-regulated under Fe-deficient conditions in wild type seedlings, while expression levels were not differing in *ttn5-1^+/-^* seedlings under both Fe supply conditions. (B), *SNX1* expression was not altered between Fe-sufficient or -deficient conditions in wild type and *ttn5-1^+/-^*seedlings. (C), *PATL2* expression did not change upon Fe supply in both wild type and *ttn5-1^+/-^*seedlings. Data were obtained from three biological replicates (n = 3). One-way ANOVA with Fisher-LSD post-hoc test was performed. Different letters indicate statistical significance (p < 0.05).

**Supplemental Table S1:** Primer list

